# SorCS2 dynamically interacts with TrkB and GluN2B to control neurotransmission and Huntington’s disease progression

**DOI:** 10.1101/2021.11.03.466767

**Authors:** Alena Salašová, Niels Sanderhoff Degn, Mikhail Paveliev, Niels Kjærgaard Madsen, Saray López Benito, Plinio Casarotto, Peter Lund Ovesen, Benedicte Vestergaard, Andreea Cornelia Udrea, Lilian Kisiswa, Lucie Woloszczuková, Islam Faress, Sadegh Nabavi, Eero Castrén, Juan Carlos Arévalo, Mai Marie Holm, Mads Fuglsang Kjølby, Ulrik Bølcho, Anders Nykjaer

## Abstract

**Background:** Huntington’s disease (HD) is a fatal neurodegenerative disorder characterized by progressive motor dysfunction and loss of medium spiny neurons (MSNs) in dorsal striatum. Brain-derived neurotrophic factor (BDNF) sustains functionality and integrity of MSNs, and thus reduced BDNF signaling is integral to the disease. Mutations in BDNF receptor SorCS2 were recently identified in HD patients. Our study investigates the role of SorCS2 in MSNs biology and in HD progression.

**Methods:** We derived a double transgenic line by crossbreeding SorCS2 deficient (KO) mice with the HD mouse model R6/1. Subsequently, we characterized the SorCS2 KO; R6/1 line by a set of behavioral and biochemical studies to evaluate phenotypes related to HD. Moreover, in combination with electrophysiology and super resolution microscopy techniques, we addressed the molecular mechanism by which SorCS2 controls synaptic activity in MSNs neurons.

**Results:** We show that SorCS2 is expressed in MSNs with reduced levels in R6/1 HD model, and that SorCS2 deficiency exacerbates the disease progression in R6/1 mice. Furthermore, we find that SorCS2 binds TrkB and the NMDA receptor subunit GluN2B, which is required to control neurotransmission in corticostriatal synapses. While BDNF stimulates SorCS2-TrkB complex formation to enable TrkB signaling, it disengages SorCS2 from GluN2B, leading to enrichment of the subunit at postsynaptic densities. Consequently, long-term potentiation (LTP) is abolished in SorCS2 deficient mice, despite increased striatal TrkB and unaltered BDNF expression. However, the addition of exogenous BDNF rescues the phenotype. Finally, GluN2B, but not GluN2A, currents are also severely impaired in the SorCS2 KO mice.

**Conclusions:** We formulate a novel molecular mechanism by which SorCS2 acts as a molecular switch. SorCS2 targets TrkB and GluN2B into postsynaptic densities to enable BDNF signaling and NMDAR dependent neurotransmission in the dorsal striatum. Remarkably, the binding between SorCS2 and TrkB or GluN2B, respectively, is mutually exclusive and controlled by BDNF. This mechanism provides an explanation why deficient SorCS2 signaling severely aggravates HD progression in mice. Moreover, we provide evidence that this finding might represent a general mechanism of SorCS2 signaling found in other brain areas, thus increasing its relevance for other neurological and psychiatric impairments.

## INTRODUCTION

Huntington’s disease (HD) is a devastating monogenic neurodegenerative disease characterized by advancing motor dysfunction, cognitive impairments, and psychiatric symptoms, which ultimately lead to the death of the patient [1, 2]. HD symptoms are caused by progressive neurodegeneration of GABAergic medium spiny neurons (MSNs) in dorsal striatum, which is followed by striatal and cortical atrophy [3, 4]. Genetically, HD patients carry an unstable CAG repeat expansion in exon 1 of the gene encoding Huntingtin protein (HTT) which translates into an abnormal N-terminal polyglutamine tract resulting in mutated Huntingtin protein (mHTT). In the healthy organism, HTT plays many important roles in neurogenesis, intracellular protein trafficking, and synapse morphology and plasticity [1, 2]. Accordingly, the accumulation of cytotoxic mHTT results in several deleterious effects such as reduced trophic stimulation and perturbed synaptic plasticity, as well as diminished synaptic delivery of N-methyl-D-aspartate (NMDA) receptors favoring their extrasynaptic accumulation. These abnormalities impair local neurotransmission while at the same time predispose MSNs to glutamate-induced excitotoxicity [2, 5–11].

Reduced levels of brain-derived neurotrophic factor (BDNF) in dorsal striatum and its impaired downstream signalling have been considered central to the altered plasticity and vulnerability of MSNs in HD [12, 13]. Through the binding to the receptor Tropomyosin related kinase receptor B (TrkB), BDNF triggers TrkB phosphorylation and activates manifold of intercellular signaling cascades that control diverse biological functions. In the mature nervous system, TrkB activation regulates neuronal survival and synaptic plasticity such as consolidation of long-term synaptic potentiation (LTP) [14–16]. For the LTP to occur, TrkB must phosphorylate the N-methyl-D-aspartate (NMDA) receptor subunits GluN2B to mediate GluN2B translocation into the synapse, and to increase the proportion of NMDA receptors that contain this subunit [17–19].

Unlike neurons in many other brain areas, striatal MSNs do not produce BDNF [20, 21] but instead depend on the BDNF release from their corticostriatal afferents [12, 22]. In mouse models of HD, reduced levels of BDNF [23] and/or TrkB [24, 25] accelerate the behavioral and pathological changes, whereas cortical overexpression of BDNF ameliorates the symptoms [26, 27]. The impaired BDNF-TrkB signalling has been mostly attributed to the reduced delivery of BDNF to the striatum due to transcriptional repression of cortical BDNF expression [5, 12], and dysfunctional anterograde axonal transport [6]. More recent studies reported that in some HD mouse models, TrkB [28] and BDNF [8] levels are normal in young symptomatic mice, but that TrkB does not properly activate the signalling pathways controlling potentiation at corticostriatal synapses. Hence, the abnormal neurotransmission is probably important at the earlier stages of HD progression, while later on, reduced trophic signalling renders the neurons vulnerable, e.g. to excitotoxicity from increased extrasynaptic glutamate stimulation.

The primary determinant of HD onset is the length of the CAG repeats on the expanded allele. Yet, this only accounts for 50-70% variations at the age of onset. These observations suggest the existence of genetic modifiers, which remain largely unknown [1, 2, 11, 29–31]. A recent genome wide association study (GWAS) of human and rat HD samples identified several single nucleotide polymorphisms (SNPs) in the Sortilin Related VPS10 Domain Containing Receptor 2 (*SORCS2)* gene [32]. SorCS2 is a member of the Vps10p-domain receptor family, so-called sortilins that are enriched in distinct neuronal populations. Receptors of this family regulate several aspects of neuronal survival, neurodevelopment, synapse remodelling, and synaptic plasticity. When dysfunctional, they are often causative to psychiatric and neurodegenerative disorders [33–36]. Conceptually, these receptors function by a dual mode of action; they can mediate intracellular sorting of ligands and co-receptors, and they can form complexes with signalling receptors at the plasma membrane to control their activity [36].

We previously discovered SorCS2 is required for BDNF-dependent plasticity in hippocampus [33]. We thus hypothesized that SorCS2 modulates BDNF-TrkB signalling in the dorsal striatum, and when absent, it could aggravate HD-related symptoms in R6/1 mice. Indeed, we found that SorCS2 deficiency severely worsens the motor symptoms present in the HD mice. Here we present a new molecular mechanism of BDNF-TrkB-SorCS2 signaling axis, and its implication in MSN plasticity, and HD progression. We show that SorCS2 enables TrkB activation and the synaptic translocation of GluN2B to MSNs synapses. While BDNF stimulates the formation of a complex between SorCS2 and TrkB to boost BDNF-TrkB signalling capacity, it uncouples SorCS2 from GluN2B resulting in enrichment of the neurotransmitter receptor in postsynaptic densities. Given that symptomatic HD mice have reduced SorCS2 levels, the attenuated activity of the receptor may be fundamental to the perturbations in synaptic plasticity that typifies HD.

## METHODS

### Mice

All mice were bred and group-housed with their littermates at Aarhus University Animal Facility with unlimited access to food and water and a 12/12 hour light/dark cycle. Generation of the SorCS2 KO mice is described in [37]. SorCS2 KO line has been backcrossed for ten generations into C57BL/6J, as were WT mice. The R6/1 mouse model (obtained from the Jackson Laboratory, USA) was originally created on a CBA/C57BL/6 hybrid background [38], but has since been backcrossed for ten generations into C57BL/6J [39]. R6/1 mice were maintained on the C57BL/6J background by crossing heterozygous males with WT females since R6/1 females are infertile. R6/1 mice were crossed into our SorCS2 KO mouse line to obtain SorCS2 KO; R6/1 line. Mice were genotyped for *Sorcs2* and HDexon1 by PCR.

### Behavioral testing

*<u>Screening for body weight, clasping and grip strength:</u>* We used WT, SorCS2 KO and R6/1 littermates as controls to SorCS2 KO; R6/1 mice. Number of mice used: WT (11 males, 10 females), SorCS2 KO (11 males, 15 females), R6/1 (12 males, 16 females), and SorCS2 KO; R6/1 mice (16 males, 8 females). Out of this cohort, two R6/1 males (both 22 weeks old) and two R6/1 females died (17 weeks and 19 weeks old). Three of the R6/1, SorCS2 KO males (two 18 weeks old and one 21 weeks old) and one SorCS2 KO; R6/1 female died (17 weeks old). Data from these death individuals are included in the graphs until the last age they were tested. The experiments were double-blinded. Mice were tested weekly for clasping and grip strength from the age of 7 weeks and until they were 22 weeks old. Body weight was monitored weekly. Clasping was tested by holding the mouse by its tail and lifting it from the top of its home-cage for 10 successive trials of three seconds, with three-second rest intervals. The testing was recorded on video, and later scored to evaluate the disease progression. Each three-second trial was scored as following: A) 0 points: Normal righting response/no certain abnormalities. Slight collapse of the hind limbs toward the midline was not scored as abnormal unless the hind limbs were at least parallel. Struggling to grab hind limbs or tail with forelimbs was not scored as abnormal. B) 1 point: One hind limb with abnormal retraction or clearly abnormal posture, or both hind limbs abnormally collapsing toward midline until they were at least parallel but without touching or crossing. C) 2 points: Both hind limbs clasping (touching or crossing) after a delay of more than one second or intermittently. D) 3 points: Almost immediate (within the first second) and persistent clasping of hind limbs. The scores from each of the 10 three second trials were summed to give a score from 0-30. Grip strength was tested by letting the mouse grip a horizontal metal bar connected to a newton meter and pulling the mouse away horizontally by the tail. The force recorded by the newton meter was stored digitally by MATLAB software. Grip strength was recorded as the highest value of 10 successive trials. For rotarod testing, open field testing and gait analysis, mice were brought to the testing room in their home-cage for at least one hour before the testing started. All equipment was wiped thoroughly with ethanol between the testing of individual mice. *<u>Rotarod testing:</u>* Rotarod Model LE 8200, Panlab. The male mice were 12 weeks old. Rotation speed accelerated gradually from 4 to 40 rpm over a 5 min period. The mice received four trials per day for three consecutive days for a total of 12 trials with no prior training, and the latency to fall was recorded for each trial. Each mouse only had one attempt to stay on the rotarod for each of the 12 trials. The mice rested in their home-cage for one hour between each trial. *<u>Open field test:</u>* Male mice were tested in the open field and with gait analysis when they were 22 weeks old. Mice were placed in the corner of an open field 40 cm x 40 cm x 35cm Plexiglas arena, and the total distance traveled during a 20 min period were recorded automatically using ANY-maze tracking software. *<u>Gait analysis:</u>* Gait analysis was performed in a homemade catwalk apparatus built by our lab. The mice walked through a 50 cm x 5.5 cm corridor with transparent Plexiglas floor to a box on the side of the apparatus containing bedding from their home cage. Green light was lit horizontally through the Plexiglas floor. Ambient red light was used to enhance contrast to the green light reflected down when the mouse placed a paw on the floor. In this way, the paw placement during walking could be accurately determined from a video recorded with the camera of an iPhone SE placed in a fixed holder under the catwalk. A custom-made MATLAB program allowed automatic calculation of stride length for each paw (distance from one placement of the paw to the next placement of the same paw) and the base with of the gait (perpendicular distance from an automatically calculated longitudinal axis of the mouse to the center of the paw). Weekly testing of clasping and grip strength was performed in the room in the animal facility where the mice were normally kept.

### Electrophysiological experiments

Male mice (P35-45) of indicated genotypes were deeply anesthetized with 4% isoflurane and decapitated. The brain was quickly extracted, transferred to ice-cold artificial cerebral spinal fluid (aCSF) bubbled with 95% O_2_ and 5% CO_2_ and cut into 400 µm coronal slices on a vibratome (Vibratome 3000 Sectioning System). *<u>Recording CA1 of hippocampus:</u>* described in [33]. *<u>Recording dorsal striatum:</u>* Slices containing striatum rostral to and including the level of the commisura anterior were transferred to a grid in a storage chamber with aCSF bubbled with 95% O_2_ and 5% CO_2,_ where they were allowed to recover for at least 1.5 hour at RT. The aCSF used for slicing and in the storage chamber contained 126 mM NaCl, 2.5 mM KCl, 1.25 mM NaH_2_PO_4_, 26 mM NaHCO_3_, 2.5 mM CaCl_2_, 1.3 mM MgCl_2_ and 10 mM D-glucose. Slices were then transferred to an interphase recording chamber perfused with aCSF bubbled with 95% O_2_ and 5% CO_2_ and containing 126 mM NaCl, 2.5 mM KCl, 1.25 mM NaH_2_PO_4_, 26 mM NaHCO_3_, 2.0 mM CaCl_2_, 1.0 mM MgCl_2_, 10 mM D-glucose and 50 µM picrotoxin (Sigma Aldrich) to block GABA_A_ receptors at a rate of 25 ml per minute. Temperature was kept at stable 31°C. Field potentials were recorded in current clamp mode using a Multiclamp 700B amplifier (Molecular Devices, Sunnyvale, CA, USA) and a glass microelectrode (7-14 MΩ) pulled on a DMZ-Universal puller (Zeitz-Instrumente, Münich Germany) filled with the same aCSF as the interphase chamber. Field excitatory postsynaptic potentials (fEPSPs) were evoked by stimulating white matter in the lower part of the corpus callosum just overlying the dorsomedial striatum with a tungsten concentric bipolar (FHC, Bowdoinham, ME, USA) electrode with pulses of 0.1 ms duration every 15 seconds before and after LTP induction. Stimulation intensity was adjusted to evoke fEPSPs at 40-50% of the maximum response on the input-output curve. 10 ng/ml BDNF (Millipore) was added to the aCSF for BDNF rescue experiments. To prevent sequestration of BDNF by unspecific binding to tube surfaces or other surfaces, albumin (0.25%) was also added to the aCSF in BDNF rescue experiments. Picrotoxin (and BDNF) was washed in for at least 20 min before starting the recording of responses. For induction of LTP we employed a theta burst stimulation (TBS) protocol adapted from [40] consisting of 10 trains spaced 15 seconds apart. Each train contained 10 bursts of four stimuli with an intraburst frequency of 50 Hz and an interburst theta frequency of 10 Hz. During the TBS protocol stimulation duration was set to 0.3 ms. LTP was induced once baseline had been stable for at least 20 min. Input-output curves were recorded by increasing stimulation intensity from 5% to 30% of 1 mA in 5% increments in the same aCSF as for LTP recordings, including 50 μM picrotoxin. To relieve the magnesium block of NMDA-Rs, all recordings of NMDA-R potentials were done in aCSF containing 0.1 mM MgCl_2_, 126 mM NaCl, 2.5 mM KCl, 1.25 mM NaH_2_PO_4_, 26 mM NaHCO_3_, 2.5 mM CaCl_2_ and 10 mM D-glucose without picrotoxin. Stimulation intensity during recording of NMDA-R potentials was fixed at 40% of 1 mA for all recordings. When the fEPSPs had been stable for at least 10 min DNQX (Sigma Aldrich), (10 µM) was washed in to block AMPA-Rs. Once the isolated NMDA-R mediated fEPSP had been stable for at least 10 min, RO-25-6981455 (Sigma) (5 µM) and PEAQX (Sigma Aldrich) (0.4 µM) was washed in to further block GluN2B- and GluN2A-containing NMDA-Rs, respectively. fEPSPs were analysed in clampfit 10 (Molecular devices) and the strength of the synaptic potential was quantified as the slope of the fEPSP. LTP was calculated as percent increase of the slope of the fEPSP compared to the average slope of the 20 min baseline. Each point on the LTP summary graphs represent the average of eight consecutive responses. E-LTP at 30 min after LTP induction was calculated as the average of the 10 minutes from 25 min to 35 min after induction of LTP for each recording. L-LTP was calculated as the average of the last 10 minutes of each recording. For analysis of experiments measuring NMDA-R potentials, the combined AMPA-R/NMDA-R response was quantified as the slope of the fEPSP as this best reflected the fast AMPA-R mediated component of the response. The slow NMDA-R component was quantified as the amplitude of the isolated NMDA-R mediated potential.

### BDNF stimulation of dorsal striatum in acute brain slices from P5 pups

P5 pups were decapitated, the brain quickly extracted and transferred to ice-cold aCSF containing 126mM NaCl, 2.5 mM KCl, 1.25 mM NaH_2_PO_4_, 26 mM NaHCO_3_, 2.5 mM CaCl_2_, 1.3 mM MgCl_2_ and 10 mM D-glucose bubbled with 95% O_2_ and 5% CO_2_ and cut into 400 µm coronal slices on a vibratome (Vibratome 3000 sectioning system). Slices containing the striatum rostral to and including the level of the commisurra anterior were transferred to a grid in a storage chamber with aCSF bubbled with 95% O_2_ and 5% CO_2_ where they were allowed to recover for at least 1.5 hour at RT. Thereafter the dorsal striatum was dissected from individual slices while they were still in oxygenated and carbonated aCSF under a stereo microscope. The dorsal striatum from matching hemi-slices was then directly transferred to aCSF continuously bubbled with 95% O_2_ and 5% CO_2_ containing either 50 ng BDNF/ml or no BDNF for 60 min at 37° C with hemi-slices from individual pups (typically three per pup) kept separated in cell strainers. Thereafter, the samples were immediately processed for the subsequent analysis.

### Perfusion, fixation, and cryosectioning of brain tissue

To perform transcardial perfusion, the mice were deeply sedated by inhaling 4% isofluorane. Per mouse, we first injected 40ml of ice-cold phosphate buffered saline (1xPBS, pH7.4) to the circulation system to remove all the blood cells. Then we perfused the mice with 40ml of ice-cold 4% formaldehyde (in 1xPBS, pH7.4) to crosslink the tissue. We removed the brain from the skull, and post-fixated it in 4% formaldehyde at +4°C for 24 hours. The next day we washed the brains four times with 1xPBS to remove the leftover formaldehyde, and then we dehydrated the brains in 30% sucrose (dissolved in 1xPBS) at +4°C until the tissue sank down. We then embedded the tissue in OCT cryopreservative and froze it to -80°C. We sectioned 14 µm thick coronal sections containing the dorsal striatum on cryostat Cryostar NX70 (Thermo Scientific), which were collected on superfrost plus slides (Thermo Scientific), and stored at -80°C until further use. All procedures were done in parallel for WT and SorCS2 KO mice.

### RNA *in situ* hybridization, immunofluorescence, and confocal microscopy

RNA *in situ* hybridization data were obtained from © 2004 Allen Institute for Brain Science. Allen Mouse Brain Atlas Available from mouse.brain-map.org/gene/show/57520. Prior the immunofluorescence, the tissue sections were thaw at room temperature for 30 min, then washed in 1xPBS (pH7.4), gently shaking for 5 min. All stainings were done in parallel for WT and SorCS2 KO sections. We performed antigen retrieval treatment using 1x target retrieval solution (DAKO, #S1699). We first preheated the solution at 96°C for 20 min in a FS3000 steamer (Braun), and only then, we added the slides, which were incubated at 96°C for 15min. The sections were transferred into 1xPBS (pH7.4) and washed three times in 1xPBS (pH7.4), gently shaking for 5 minutes. Afterwards, the sections were blocked in PBTA solution (0.1% BSA, 0.3% TritonX-100 in PBS, pH7.4) containing 5% donkey serum at room temperature for 1h. The sections were incubated with primary antibodies (diluted in PBTA-5% donkey serum) in a wet chamber overnight. We used following primary antibodies and dilutions: anti-murine-SorCS2 1:100 (#AF4237, R&D Systems), anti-DARPP-32 1:100 (#AB40801, Abcam), anti-NeuN 1:100 (#MAB377, Millipore), anti-GFAP 1:500 (#Z0334, DAKO), anti-parvalbumin 1:500 (#AB11427, Abcam), Anti-calretinin 1:500 (#AB5054, Merck Millipore), anti-NPY 1:100 (#11976, Cell Signaling Technology), anti-ChAT 1:100 (#ab70219, Abcam). The next day, the sections were washed 4 times x 5min in 1xPBS (pH7.4) at room temperature, gently shaking. Sections were then incubated with the secondary antibodies (1:800) in PBTA-5% donkey serum for 1.5h at room temperature. We also added Hoechst 1:2000 to stain nuclei. We used AlexaFluor-488 and AlexaFluor-568 conjugated secondary antibodies (Molecular Probes, Invitrogen) of suitable species reactivity. Labelled sections were mounted in Fluorescence Mounting Medium (#S3023, DAKO) and covered by long coverslips. Dorsal striatum was imaged with Zeiss LSM780 or LSM800 confocal microscopes using 10X/0.45, and 40x or 63x/1.20 objectives. We first adjusted the settings to the SorCS2 KO tissue and then we applied the same settings for the WT. Acquired images were post-processed in Adobe Photoshop and Adobe Illustrator.

### Proximity Ligation Assay (PLA) and super-resolution microscopy

The hemi-slices were handled as free-floating sections. First, they were fixed in 4% formaldehyde for 1 hour, washed 3 times with 1xPBS pH7.4 containing 0.1% TritonX-100. Then they were blocked in PBTA solution (described above) for 1 hour in room temperature. For the GluN2B-PSD95 immunofluorescence staining, these primary antibodies were used: anti-GluN2B (1:200, #ab65875, Abcam) and anti-PSD95 (1:200, #ab2723, Abcam). The slices were incubated with primary antibodies overnight at +4C. Next day, the slices were washed 3 times with 1xPBS pH7.4 containing 0.1% TritonX-100. We then incubated the slices in secondary donkey antibodies conjugated to AlexaFluor488 and AlexaFluor568 (Molecular Probes, Invitrogen) for 2 hours at room temperature. After that, the slices were washed 3 times with 1xPBS pH7.4 containing 0.1% TritonX-100. Then they were placed on Superfrost plus slides (Thermo Scientific), mounted with Fluorescence Mounting Medium (#S3023, DAKO) and covered by long glass coverslips. For the Sorcs2-GluN2B (PLA) + PSD95 (immunofluorescence) staining, the hemi-slices were blocked as described above and incubated with primary antibodies anti-GluN2B (1:200, #ab65875, Abcam), anti-PSD95 (1:200, #ab2723, Abcam), and anti-murine-SorCS2 (1:250, #AF4237, R&D Systems) at 4°C overnight. The next day, the slices were washed 3 times with 1xPBS pH7.4 containing 0.1% TritonX-100. Then we performed the Duolink Fluorescent PLA protocol using Duolink Prober Maker Kit (DUO96010, Sigma Aldrich) as recommended by the manufacturer. We used anti-goat plus (DUO92003, Sigma Aldrich) and anti-rabbit minus (DUO92005, Sigma Aldrich) Duolink PLA probes for the detection of the GluN2B-SorCS2 in close proximity. Secondary donkey anti-mouse antibodies conjugated to AlexaFluor488 (for the detection of PSD95) were applied for 2 hours at room temperature in combination with the far-red detection PLA staining according to the manufacturer protocol. Dorsal striatum was imaged with Zeiss LSM800 confocal microscope using the Airyscan mode for the super-resolution imaging, x63 objective, and voxel size 40×200nm. Co-localization of GluN2B to PSD95 was quantified with the FIJI software as the object area positive for both GluN2B and PSD95 divided by the total object area positive for GluN2B. Co-localization of PSD95 to the SorCS2-GluN2B PLA signal was quantified with the FIJI software as the object area positive for both the PLA and PSD95 divided by the total object area positive for PSD95. All intensity thresholds were obtained with the Fiji autothreshold command. All pixels of the GluN2B-, PLA- and PSD95-positive objects were quantified via the AnalyzeParticles Fiji command. The 3D reconstruction was performed by Imaris 8.2 software.

### Tissue dissection and protein extraction

Mice were sacrificed by cervical dislocation. The brain was quickly removed, and transferred to ice-cold 1xPBS, pH7.4. The dorsal striatum (upper two thirds of the striatum) rostral to and including the commisurra anterior, and the overlying cortex was dissected out in 1xPBS, pH7.4. The tissue was immediately frozen on dry ice and stored at – 80°C. For western blots, the tissue was homogenized in ice-cold lysis buffer (10mM Tris-base, 150mM NaCl, 1mM EDTA, 1% NP40, pH8) containing freshly added 1x proteases inhibitors (#88666, Thermo Scientific) and 1x phosphatases inhibitors (#4906837001, Roche). In case of stimulated striatal hemi-slices, we immediately homogenised them in separate tubes containing 200 uL ice cold lysis buffer (10mM Tris-base, 150mM NaCl, 1mM EDTA, 1% NP40, pH8) containing freshly added 2x proteases inhibitors (#88666, Thermo Scientific) and 2x phosphatases inhibitors (#4906837001, Roche), and 4.5 nM sodium ortovanadate. Samples were centrifuged at 1412g at +4°C for 15 min before the supernatant was transferred to a new tube and kept at - 80° C until further analysis.

### Real time quantitative PCR (qPCR)

RNA was purified using the Nucleospin gel and PCR clean-up kit from Macherey-Nagel. Complementary DNA (cDNA) was synthetized by using High-Capacity RNA-to-cDNA Kit (#4387406, Applied Biosystems) according to the manufacturer’s protocol. For qPCR reaction, TaqMan Fast Advanced Master Mix (#4444556, Applied Biosystems) and TaqMan probes (GAPDH Mm99999915_g1, SorCS2 Mm00473050_m1; Thermo Fisher Scientific) were used according to manufactureŕs instructions. The reaction was performed in QuantStudio 7 Flex Real-Time PCR System (Applied Biosystems) using the following program: an activation step at 95°C for 20s, followed by 40 cycles of denaturation at 95°C for 1s and annealing at 60°C for 20s. Next, ΔCp values for each of the samples were calculated by using GAPDH as the reference gene.

### ELISA

*<u>BDNF ELISA:</u>* Dorsal striatum and overlying cortex was dissected from 22-week-old mice. The tissue was lysed in ice-cold lysis buffer (10 mM Tris pH 7.4, 150 mM NaCl, 2 mM EDTA, 1% NP-40, 1% SDS, 1 mM PMSF, 1 μg/ml Aprotinin, 2 μg/ml Leueptine, 1 mM Vanadate, 10 mM NaF and 20 mM β-glycerophosphate) and centrifuged at 14.000 x g for 15 min. For each sample BDNF concentration detected by ELISA was normalized to total protein in the lysate determined by the Bradford method. Detailed protocol is described in [41]. *<u>TrkB-SorCS2 ELISA:</u>* Hippocampal primary cell culture was prepared as described previously [42] from E18 rat embryos. The cells were seeded on 24-well plate and kept in the culture for DIV8-10. Following the treatment with BDNF, cells were lysed for 30min shaking in NP lysis buffer (3M Tris-HCl pH8, 5M NaCl, 0.5M NaF, 1% NP40, 10% glycerol). The ELISA was performed according to the published protocol [43] using these antibodies: anti-TrkB goat 1:1000 (#AF1494, R&D systems), anti-SorCS2 rabbit 1:2000 (#bs-11963R BIOSS), anti-Rabbit IgG HRP-conjugated antibody 1:5000 (#170-5046 BioRad). First, 96-well plate (OptiPlate 96, Perkin Elmer) was coated with anti-TrkB antibody (1:1000 in carbonate buffer pH9.8) at +4°C overnight. Then we blocked the plate with 2% BSA in 1xPBS pH7.4 with 0.1% Tween20 for 2h at RT, shaking. Afterwards, we added the lysates (120ug of total protein, 100ul/well), sealed the plate, and incubated on a shaker overnight at +4°C. Next, we washed the plate 3 times with 1xPBS pH7.4 with 0.1% Tween20, and added anti-SorCS2 antibody (1:2000 in 2% BSA in 1xPBS pH7.4 with 0.1% Tween20). We sealed the plate and incubated it on a shaker for 2h at RT or at +4°C overnight. On the third day we washed the plate 3 times with 1xPBS pH7.4 with 0.1% Tween20 and added anti-rabbit IgG HRP-conjugated antibody (RRID:AB_11125757, 100ul/well). We sealed the plate and incubated it on a shaker at RT for 2h. In the end, we washed the plate 3 times with 1xPBS pH7.4 with 0.1% Tween20, and developed the luminescence using premixed ECL reagent (1:1, Pierce, 100ul/well).

### Co-immunoprecipitation, Synaptosome extraction, and Western blotting

*<u>Co-immunoprecipitation:</u>* GammaBind G Sepharose beads (50ul/sample; #17-0885-01, GE Healthcare) were washed in 1xPBS (pH7.4) and coated with 1ug of anti-TrkB antibody (#AF1496, R&D Systems) or 1ug of normal Rabbit IgG (#AB-105-C, R&D Systems) while rotating in 1xPBS (pH7.4) ON at 4 ͦC. The beads were then washed 3 times in 1xPBS (pH7.4), and incubated with 500ul of the lysates of dorsal striatum while rotating at +4°C overnight. The next day, the beads were washed 5 times in 1xPBS-pH7.4 containing 0.05% Tween-20. SDS sample buffer (#NP0007, Thermo Fisher Scientific) and 20mM DTT was then added to the washed beads, proteins were denatured by heating to 95°C for 5 min and subjected the SDS-page. *<u>Synaptosome preparation:</u>* For the preparation of synaptosome fraction, we used Syn-PER synaptic protein extraction reagent (#87793, Thermo Scientific). For the extraction, we followed the protocol provided by the manufacturer. In the end of the extraction, we added 1ml of the Syn-PER reagent per 1g of tissue. We resuspended the pellets and measure the protein concentration. We loaded 30ug of protein per well into gradient 4-12% NuPAGE Bis-Tris acryl amid gels (#NP0321, Thermo Fisher Scientific) and performed western blotting analysis.*<u>Western blotting:</u>* Protein concentration of lysates was determined with Bicinchoninic Acid Kit (#BCA1, Sigma Aldrich). Equal amount of proteins was loaded on the polyacrylamide gels, separated by SDS-page, and transferred to a PVDF membrane by Western blotting in 25 mM Tris-base and 192 mM glycine (pH 8.0). Membranes were blocked in 50 mM Tris-base, 0.5 M NaCl, 2% skimmed milk powder and 2.1% Tween-20. After blocking membranes were washed for 2 x 5 min in washing buffer containing; 10 mM HEPES, 140 mM NaCl, 2 mM CaCl_2_, 1 mM MgCl_2_, 0.2% skimmed milk powder and 0.05% Tween-20 (pH 7.8). Membranes were then incubated with primary antibodies diluted in washing buffer ON at 4 C. Next day membranes were washed for 3 x 5 min and incubated with HRP-coupled secondary antibodies (DAKO) diluted 1:3000 in washing buffer for 1.5 hour at RT. After washing again for 5 x 5 min western blots were visualized with ECL plus Western Blotting detection system (#RPN2231, GE Healthcare) or Immobilon Forte Western HRP substrate (#WBLUF, Sigma Aldrich) dependent on the signal. We developed the membranes on developing machine LAS4000 (Fuji film). We used following primary antibodies and dilutions: anti-(murine)SorCS2 1:1000 (#AF4237, R&D Systems), anti-DARPP-32 1:1000 (#AB40801, Abcam), anti-BDNF (#P23560, Icosagen), anti-TrkB 1:1000 (#AF1494, R&D Systems), anti-phospho TrkA (Tyr674/675)/TrkB (Tyr706/707) (C50F3) 1:1000 (#4621, Cell signalling technology), anti-GluN1 1:1000 (#5704, Cell signalling technology), anti-GluN2A 1:1000 (#4205, Cell signalling technology), anti-GluN2B 1:1000 (#06-600, Sigma Aldrich), anti-PSD95 1:1000 (#3450, Cell signalling technology), anti-Synaptophysin 1:1000 (#5461T, Cell signalling technology), anti-ERK1/2 1:2000 (#4695, Cell signalling technology), anti-pERK1/2 1:1000 (#4370, Cell signalling technology), anti-pAkt 1:2000 (#4060, Cell signalling technology), anti-beta-Actin 1:5000 (#A5441, Sigma). The immunoblots were quantified by densitometry in Multigage V3.0 software (Fuji film) or ImageJ software. We substracted background and normalized the signal to loading control. For BDNF stimulation assay, we normalized phosphorylated TrkB to total levels of TrkB.

### Statistics

We used GraphPad Prism 9.1.2 for the data visualization and statistical analysis. All data sets were evaluated for normality and statistical outliers. The data are presented as mean ±SEM. For simplicity, the symbols * indicate p<0.05, ** indicate p<0.01, *** indicate p<0.001, and **** indicate p<0.0001. An overview of used statistical tests can be found in the **Table 1**.

**Table 1:**
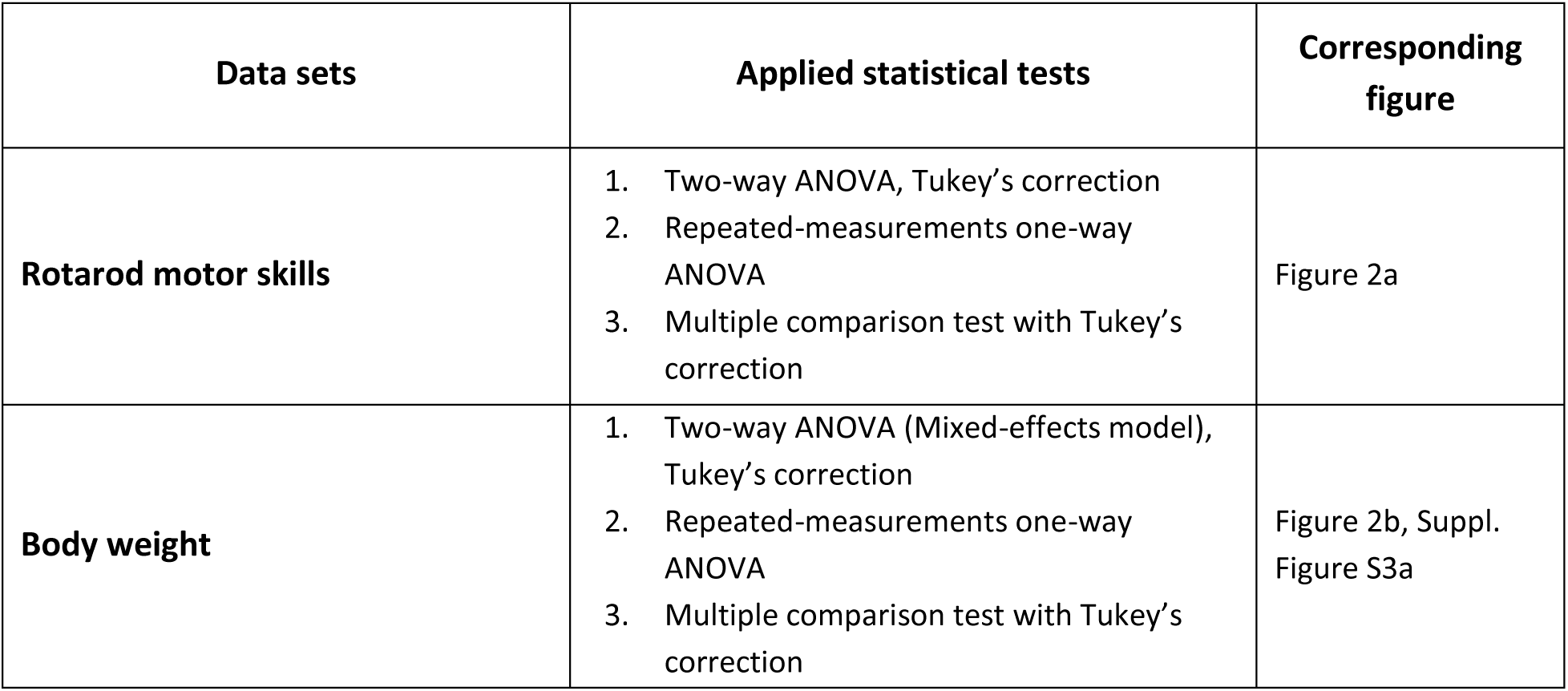
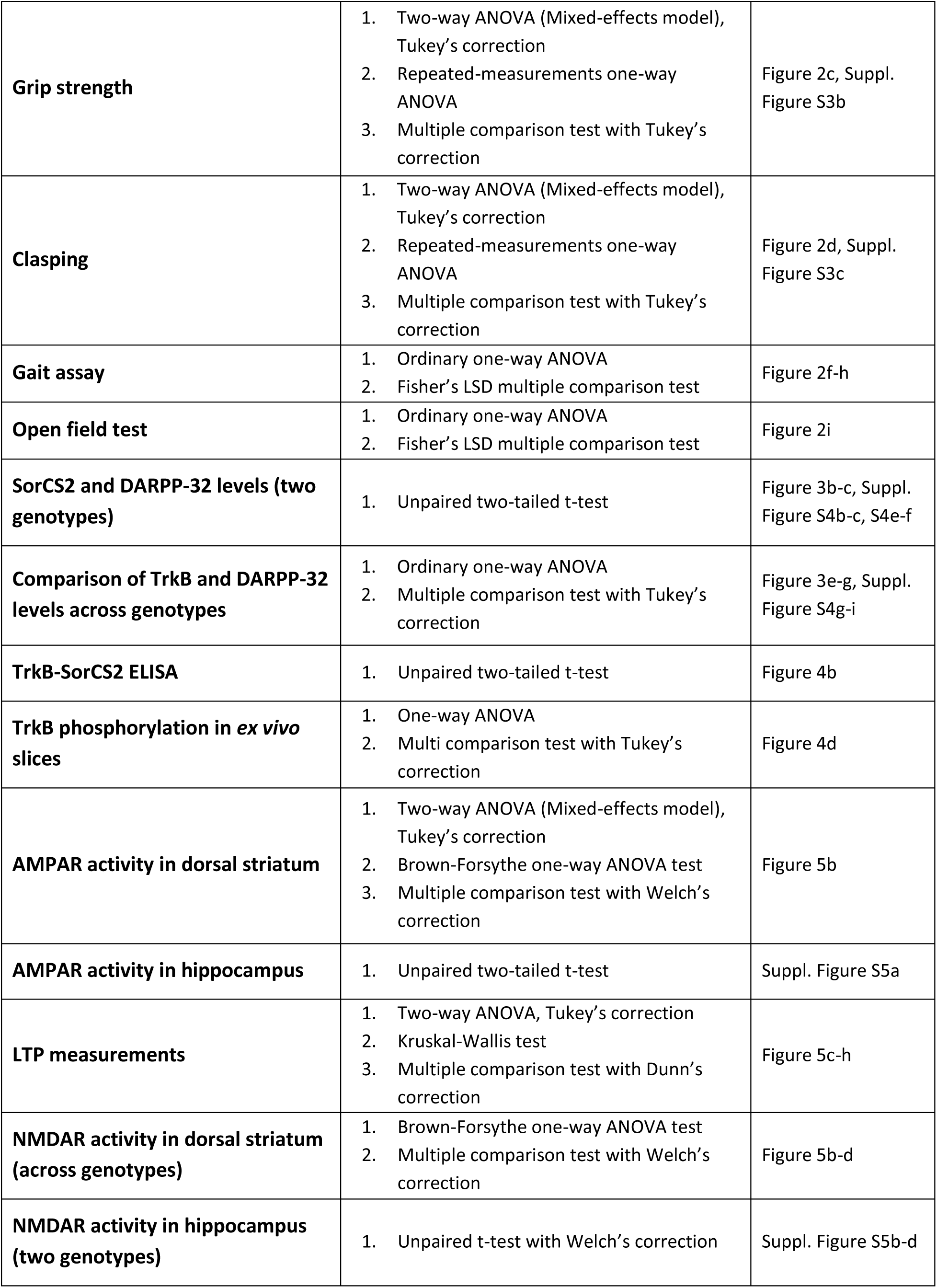
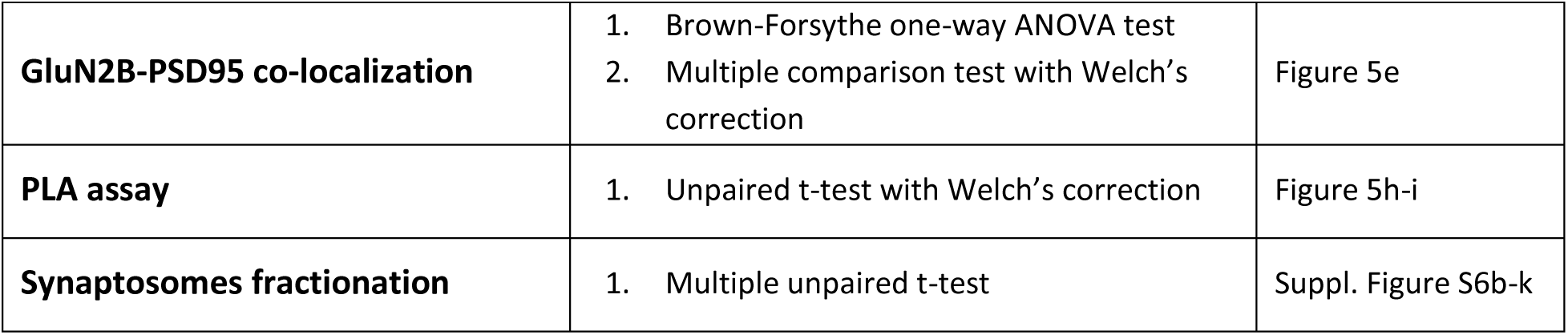
Overview of the statistical testing used in the study

## RESULTS

### SorCS2 is selectively expressed by MSNs in dorsal striatum

SorCS2 is typically expressed by neurons in the adult mouse brain, with the highest expression in dentate gyrus of the hippocampus and piriform cortex [33, 44]. To explore SorCS2 expression profile in corticostriatal circuits, we first took advantage of the available *in situ* hybridization data (IHC) from Allen brain atlas. We observed that besides the piriform cortex, SorCS2 is highly expressed in dorsal striatum (caudoputamen), cortical layers II and III (including motor cortex and anterior cingulate cortex), induseum griseum, and intermediate lateral septal nucleus (**Figure 1a-c**). To determine which cell types express SorCS2 in dorsal striatum, we performed immunofluorescence using a number of known cell type-specific markers. Age-matched SorCS2 knockout (KO) brain tissue served as negative control. We found that SorCS2 is expressed by DARPP-32 and NeuN positive neurons (**Figure 1d-e**) that designates MSNs in dorsal striatum. On contrary, SorCS2 was absent from any of the most prominent types of striatal interneurons [45, 46] such as calretinin positive interneurons, large aspiny cholinergic interneurons (ChaT+), GABAergic fast-spiking parvalbumine-positive interneurons, and low-threshold-spiking neuropeptide Y (NPY+) interneurons (**Suppl. Figure S1a-d**). As previously reported for the hippocampus, SorCS2 was absent in Glial fibrillary acidic protein positive (GFAP+) glia cells in dorsal striatum (**Suppl. Figure S2**). Our data suggest that SorCS2 may be involved in the biology of MSNs in the dorsal striatum.

**Figure 1:**
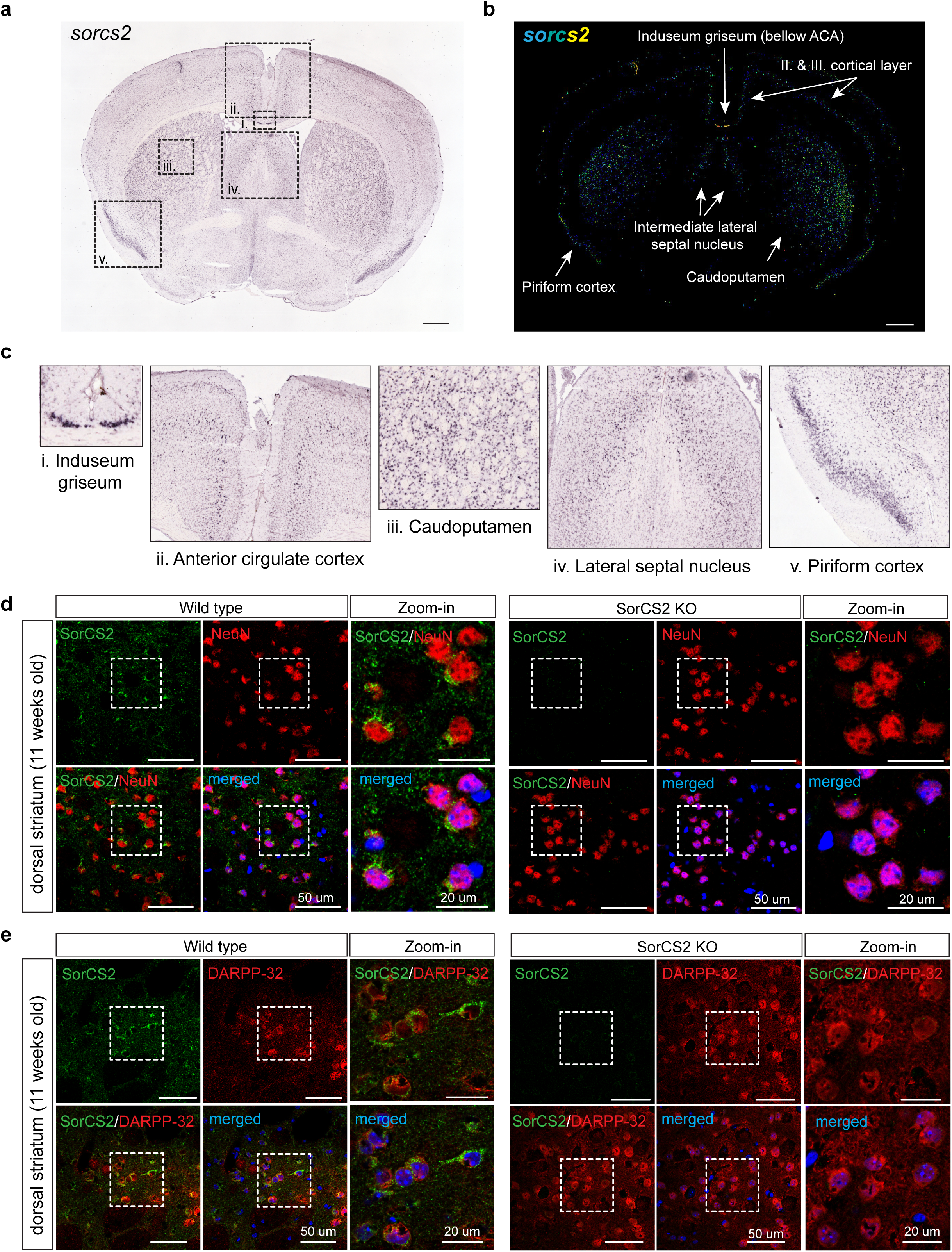
SorCS2 is expressed by MSNs in corticostriatal circuits. **a** RNA *In situ* hybridization shows *sorcs2* expression in coronal section of P56 WT mouse brain, including the corticostriatal circuits. **b** Quantified expression profile of *sorcs2* from the same section. Scale-bar is 700um. **c** *Sorcs2* is expressed in multiple brain regions including induseum griseum (i.), interior cirgulate cortex (ii.), caudoputamen (iii.), lateral septum (iv.), and piriform cortex (v.). **a-c** Image credit: ©2004 Allen Institute for Brain Science. Allen Mouse Brain Atlas Available from mouse.brain-map.org/gene/show/57520. **d-e** Immunofluorescence followed by confocal microscopy imaging revealed that SorCS2 is expressed by NeuN (**d**) and DARPP-32 (**e**) positive MSNs neurons in dorsal striatum of adult mice. Blue channel corresponds to nuclei stained by Hoechst. The scale bar corresponds to 50µm, and to 20µm in magnified figures.

### SorCS2 deficiency accelerates HD progression in R6/1 mice

Since MSNs in dorsal striatum control motor skills, we studied motor function in SorCS2 KO and tested whether SorCS2 deficiency worsens HD-related motor symptoms. To address this, we generated double transgenic line by crossing SorCS2 KO mice into the R6/1 transgenic HD model, which carries a human 115 CAG repeat in exon 1. First, we subjected 12 weeks old males to rotarod behavior testing to study motor skill learning. Mice were placed on the accelerating rotarod without any prior training and received four trials per day for three consecutive days. Each mouse had a single attempt on the rotarod for each trial and we recorded the latency to fall (**Figure 2a**). This experiment revealed a significant role of genotype (p=0.014) and trial number (p=<0.0001) in motor performance. SorCS2 KO mice and R6/1 mice both displayed impaired motor skill learning, as did the double transgenic SorCS2 KO; R6/1 mice [SorCS2 KO vs WT (p=0.0006), R6/1 vs WT (p=<0.0001), SorCS2 KO; R6/1 vs WT (p=0.0002), SorCS2 KO vs R6/1 (p=0.0154)]. At this age, there was no additive effect of SorCS2 loss of function to motor learning and HD progression in the R6/1 line.

**Figure 2:**
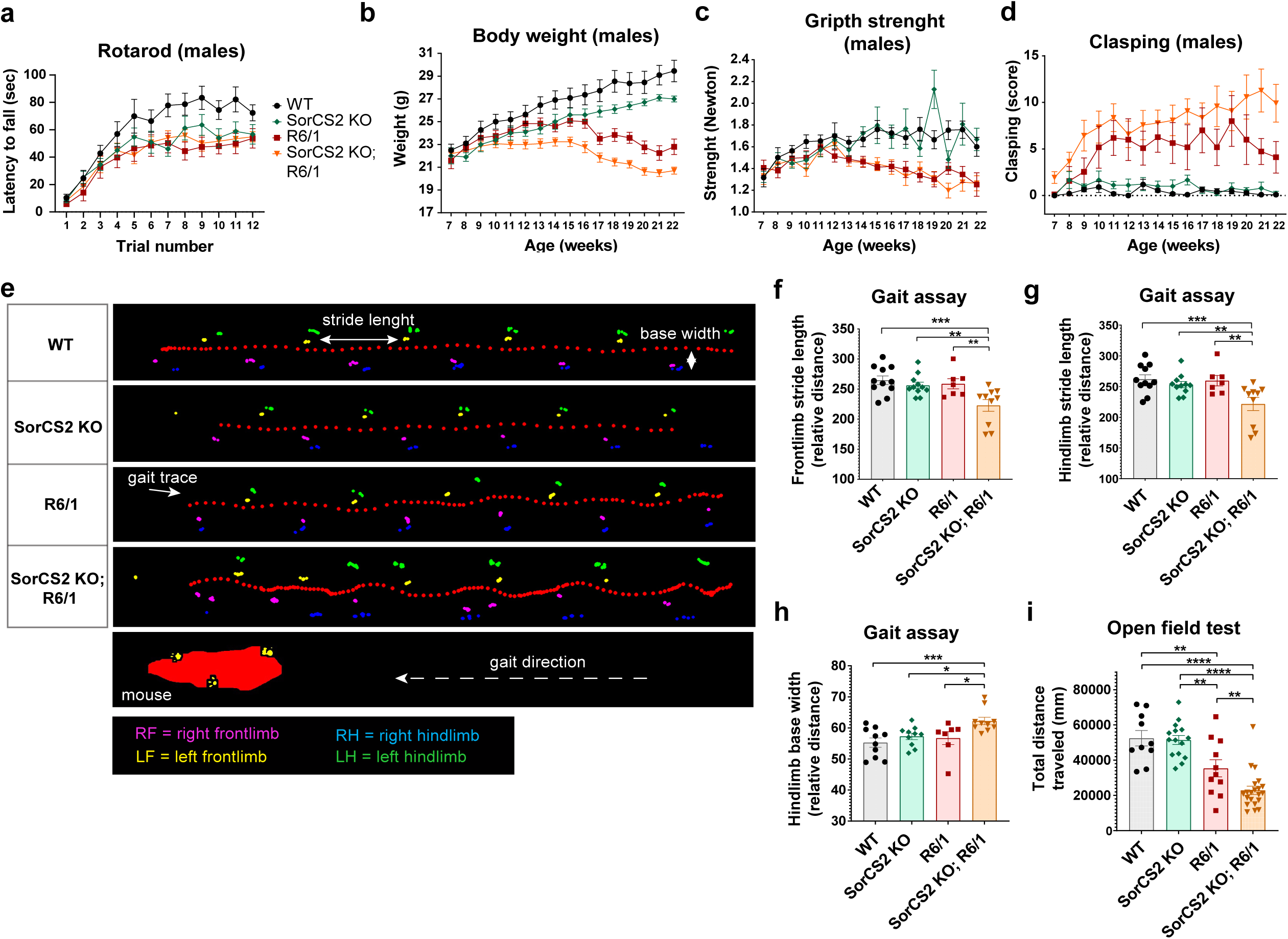
SorCS2 accelerates and worsens HD disease progression. **a** Rotarod motor skill learning in 12-weeks-old males revealed contribution of genotype (p=0.014) and the trial number (p<0.0001) (two-way ANOVA, WT: n=9, SorCS2 KO n=10). Repeated measurements one-way ANOVA and Tukey’s multiple comparisons test (MC): SorCS2 KO vs WT (p=0.0006), R6/1 vs WT (p<0.0001), SorCS2 KO; R6/1 vs WT (p=0.0002), SorCS2 KO vs R6/1 (p=0.0154); **b-d** Staging of animals over time. WT; n=11, SorCS2 KO: n=10, R6/1: n=12, SorCS2 KO;R6/1: n=13. **b** Body weight with contribution of genotype and age (both p<0.0001, two-way ANOVA). MC: SorCS2 KO vs WT (p<0.0001), R6/1 vs WT males (p=0.0006), SorCS2 KO; R6/1 vs R6/1 (p<0.0001), SorCS2 KO; R6/1 vs SorCS2 KO (p=0.0028). **c** Grip strength with contribution of genotype (p=<0.0001) and age (p=0.0004) (two-way ANOVA). MC: R6/1 vs WT (p<0.0001). **d** Clasping score revealed contribution of genotype (p<0.0001), age (p=0.0096), and genotype x age (p=0.0118) (two-way ANOVA). MC: SorCS2 KO vs WT (p=0.0059), R6/1 vs WT (p<0.0001), SorCS2 KO; R6/1 vs WT (p<0.0001), SorCS2 KO; R6/1 vs R6/1 (p<0.0001). **e** Representative visualization of gait patterns across genotypes. **f** Quantification of the front limb stride length; p=0.0017 for difference among genotypes (one-way ANOVA), MC: WT vs SorCS2 KO; R6/1 (p=0.0003), SorCS2 KO vs SorCS2 KO; R6/1 (p=0.003), R6/1 vs SorCS2 KO; R6/1 (0.0043) (Fisher’s LSD test). **g** Hind limb stride length; p=0.003 for difference among genotypes (one-way ANOVA), MC: WT vs SorCS2 KO; R6/1 (p=0.0007), SorCS2 KO vs SorCS2 KO; R6/1 (p=0.0061), R6/1 vs SorCS2 KO; R6/1 (p=0.0038) (Fisher’s LSD test). **h** Hind limb base width; p=0.0061 for difference among genotypes (one-way ANOVA). MC: WT vs SorCS2 KO; R6/1 (p=0.0009), SorCS2 KO vs SorCS2 KO; R6/1 (p=0.0141), R6/1 vs SorCS2 KO; R6/1 (p=0.0133) (Fisher’s LSD test). **i** Total distance traveled of 22-weeks-old males in the open field test revealed significant contribution of the genotype (p<0.0001) (one-way ANOVA). MC: WT vs R6/1 (p=0.0025), WT vs SorCS2 KO; R6/1 (p<0.0001), SorCS2 KO vs R6/1 (p=0.0018), SorCS2 KO vs SorCS2 KO; R6/1 (p p<0.0001), R6/1 vs SorCS2 KO; R6/1 (p=0.009) (Fisher’s LSD test). The graphs show mean values ±SEM.

To study the progression of HD, we next monitored the mice from 7 weeks to 22 weeks of age with respect to body weight, grip strength, and clasping. Both, SorCS2 KO and R6/1 males had lower body weight than WT (p<0.0001 and p=0.0006, respectively), which was further reduced in SorCS2 KO; R6/1 mice when compared to R6/1 (p=<0.0001) and SorCS2 KO (p=0.0028) (**Figure 2b**). The same applied to females [SorCS2 KO vs WT (p=<0.0001), R6/1 vs WT (p=<0.0001), SorCS2 KO; R6/1 vs R6/1 (p=<0.0001)] (**Suppl. Figure S3a**). Grip strength was tested by letting the mouse grip a horizontal metal bar connected to a newton meter, and pulling the mouse away horizontally by the tail. Grip strength was diminished in R6/1 mice in both males (p=<0.0001) (**Figure 2c**) and females (p=0.0009) (**Suppl. Figure S3b**). However, SorCS2 KO mice did not stand out from WT, and neither did receptor deficiency aggravate the phenotype of R6/1 in this paradigm. We next tested for clasping, which is a marker of neurodegeneration commonly used to study disease progression in HD mice [38]. The mice may initially present a dystonic posture that, as the disease builds up, eventually develops into overt clasping of the hind limbs when lifted by the tail. SorCS2 KO displayed minor clasping when compared to WT in both males (p=0.0059) (**Figure 2d**) and females (p<0.0001) (**Suppl. Figure S3c**). As expected, R6/1 mice exhibited significant clasping [(p<0.0001), both genders]. Strikingly, we observed a severe worsening of clasping in SorCS2 KO; R6/1 mice when compared to R6/1 [(p<0.0001) both genders], which accelerated with aging (males: p=0.0118, females: p<0.0001). This observation manifested an important contribution of SorCS2 loss of function on HD progression.

To assess gait performance, we recorded the movement of 22 weeks old males using an in-house-built catwalk apparatus. The assay enabled us to distinguish single steps of a walking mouse, and measure the stride length and the base width of the gait (**Figure 2e, Suppl. Figure S3d**). Representative videos for WT, SorCS2 KO, R6/1, and SorCS2 KO; R6/1, respectively, are available in **Suppl. Video SV1-4**). The double transgenic line showed impaired gait pattern through the corridor (**Figure 2e, Suppl. Figure S3e**). Image analysis revealed that SorCS2KO; R6/1 mice have significantly reduced front limb stride length compared to the R6/1 (p=0.0043), while the single transgenic lines performed as WT (**Figure 2f**). Hind limb stride length was also significantly reduced in SorCS2 KO; R6/1 mice compared to R6/1 mice (p=0.0038) (**Figure 2g**), and hind limb base width was increased only in SorCS2KO; R6/1 compared to the other genotypes; to WT (p=0.0009), SorCS2 KO (p=0.0141), and R6/1 (p=0.0133) (**Figure 2h**). These data indicate that the impaired gait performance is caused by cumulative effect of SorCS2 loss of function in the HD model. Finally, we subjected 22 weeks old males to the open field test. In this test, R6/1 as well as SorCS2KO; R6/1 mice moved less than WT (p=0.0061 and p<0.0001, respectively) and with significant worsening of the activity in SorCS2 KO; R6/1 compared to the R6/1 mice (p=0.0082) (**Figure 2i**). Taken together, our data demonstrate that SorCS2 loss of function accelerates and worsens some of the HD-related motor symptoms in R6/1 mice.

### Striatal expression of SorCS2 and DARPP-32 during HD progression

To investigate whether HD progression affects SorCS2 expression, we examined RNA and protein levels in our transgenic mice using the real-time quantitative PCR (qPCR) and western blot analyses. While SorCS2 expression in the dorsal striatum of 6 weeks old asymptomatic R6/1 mice did not differ from WT mice (**Suppl. Figure S4a-b**), it was reduced by 50% both at protein (p=0.0001) and RNA levels (p=0.0019) in 22 weeks old symptomatic animals (**Figure 3a-c**). This difference was tissue specific as SorCS2 expression in cortex was unaltered (**Suppl. Figure S4d-f**). Hence, a reduction of striatal SorCS2 expression may accelerate and exacerbate the HD progression. A decline in DARPP-32 is known to reflect the neurodegenerative process of MSNs in several HD mouse models [23, 47]. We found that DARPP-32 expression in 6 weeks old asymptomatic R6/1 mice was normal (**Suppl. Figure S4a, c**). However, at 22 weeks of age, DARPP-32 was significantly reduced in both R6/1 (p=0.0082) and SorCS2 KO; R6/1 mice (p=0.0044). DARPP-32 was unaltered in *Sorcs2^−/−^* mice of similar age (**Figure 3d, 3g**). The data suggest that a secondary reduction in SorCS2 expression may fuel the HD progression, possibly by altering MSNs function rather than by directly affecting their degeneration.

**Figure 3:**
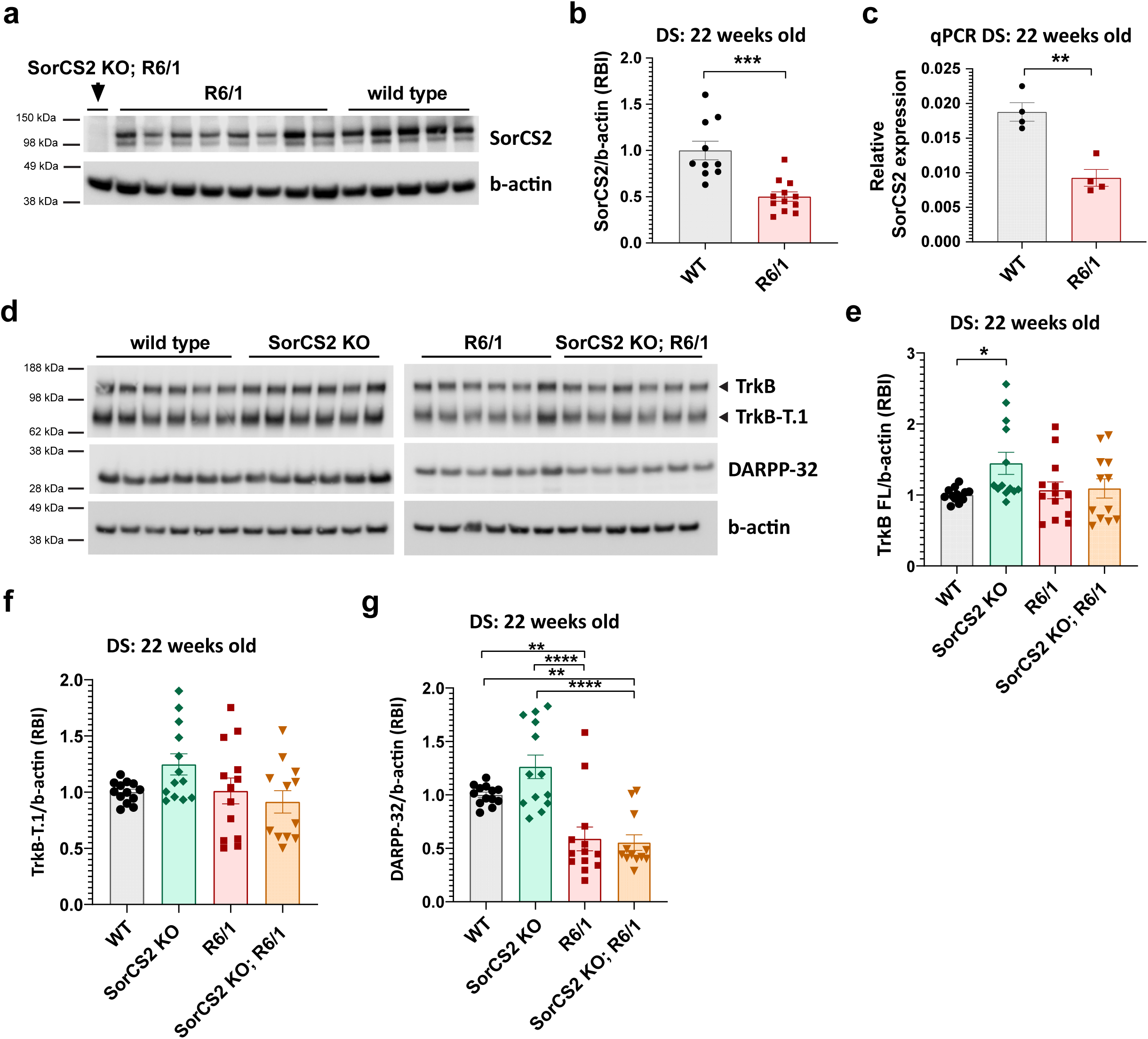
Analysis of DARPP-32, TrkB, and SorCS2 protein levels in dorsal striatum of 22-weeks old males. **a** Immunoblot of SorCS2 protein levels in dorsal striatum of 22-weeks old males, quantified in (**b**) show depleted levels of SorCS2 in R6/1 mice (p=0.0001). **c** Real-time qPCR analysis of SorCS2 expression in dorsal striatum confirmed decreased levels of SorCS2 in R6/1 mice (p=0.0019); Unpaired two-tailed t-test (**b-c**). **d** Representative immunoblots show detection of DARPP-32, the full-length (FL) and truncated (T.1) TrkB, and b-actin as loading control. Quantification of the immunoblots followed by ordinary one-way ANOVA and Tukey’s multiple comparisons test revealed that SorCS2 KO have increased levels of FL-TrkB (p=0.0482) (**e**) but there were no changes in its truncated form (**f**). **g** Quantification of DARPP-32 show significant differences dependent on the genotype: R6/1 vs WT (p=0.0082), SorCS2 KO; R6/1 vs WT (p=0.0044), SorCS2 KO vs R6/1 (p<0.0001), SorCS2 KO; R6/1 vs SorCS2 KO (p<0.0001); Ordinary one-way ANOVA and Tukey’s multiple comparisons test (**e-g**). RBI = Relative band intensity. Graphs show mean values ±SEM.

### SorCS2 is required for robust TrkB signaling in dorsal striatum

Reduced BDNF signaling is fundamental to the neurodegenerative process and altered neurotransmission of MSNs. In some HD mouse models, the cortical transcription of BDNF, its anterograde transport and release from cortical terminals in the dorsal striatum, as well as the TrkB expression are suppressed [6, 7, 12, 13, 23, 48]. In other cases, BDNF and TrkB expression is preserved but TrkB signaling is compromised [8, 28, 41]. We therefore quantified BDNF and TrkB levels in corticostriatal circuits to study the impact of SorCS2 on TrkB signaling. Despite severe motor symptoms in the R6/1 and SorCS2 KO; R6/1 mice, BDNF levels were not reduced in cortex nor in dorsal striatum, indicating that the production and trafficking of BDNF is unaffected by mHTT and SorCS2 (**Suppl. Figure S4g-j**). However, we observed an increase in BDNF levels in dorsal striatum of SorCS2 KO; R6/1 when compared to SorCS2 KO (**Suppl. Figure S4h)** which might indicate a compensation in BDNF secretion to overcome the neurodegeneration. There was also no difference in BDNF maturation, as we observed no shifts in the ratio between proBDNF and BDNF levels in dorsal striatum (**Suppl. Figure S4i-j**). Expression of the full-length TrkB receptor was slightly increased in the SorCS2 KO mice (p=0.0482) but this difference disappeared on the R6/1 background (**Figure 3d-e**). The expression of the truncated TrkB variant (named TrkB-T.1) was independent of genotype (**Figure 3d, f**). We next investigated whether SorCS2 may physically interact with TrkB to control its signalling abilities. To this end, TrkB was immunoprecipitated from dorsal striatum and the precipitate probed for SorCS2. Indeed, we found that the two receptors co-immunoprecipitated demonstrating that SorCS2 and TrkB can physically interact to form a heteromeric complex (**Figure 4a**). Strikingly, in primary hippocampal cultures, this interaction was increased by approximately 200% when treated with 10 ng/ml BDNF (**Figure 4b**). We then asked if SorCS2 enables BDNF-TrkB signaling in the histologically intact dorsal striatum. BDNF-induced TrkB phosphorylation in *ex vivo* striatal slices is detectable in newborn pups, peaks around P7, thereafter it declines, and disappears around P12 [49, 50]. Hence, we prepared acute brain slices from P5 pups that were maintained in artificial cerebral spinal fluid (aCSF). Dorsal striatum from one set of hemi-slices was stimulated with 50 ng/ml BDNF (in aCSF) for 1 hour, while the other set of matching hemi-slices were incubated only in aCSF serving as unstimulated control. BDNF treatment induced a 2.3-fold increased phosphorylation of TrkB when normalized to total TrkB (p=0.0023) (**Figure 4c-d**). Strikingly, when comparing the response to BDNF treatment in WT and SorCS2 KO mice, we found that TrkB phosphorylation was reduced by 63% in slices from the knockouts (p=0.0047) (**Figures 4d**). We thus conclude that SorCS2 binds TrkB in a BDNF-dependent manner, and that this interaction is required for robust TrkB phosphorylation in the dorsal striatum.

**Figure 4:**
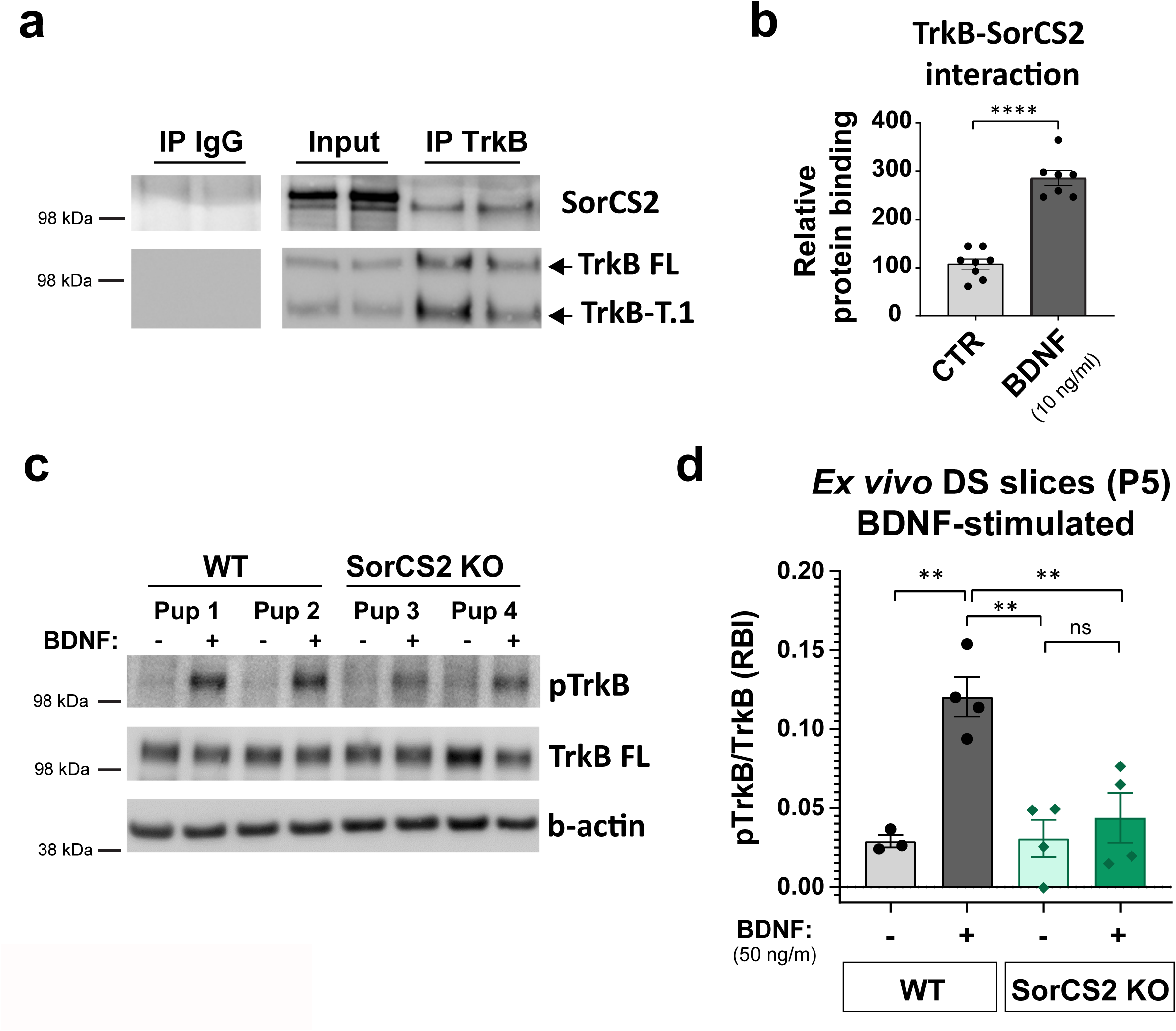
SorCS2 interaction with TrkB in dorsal striatum is required for TrkB phosphorylation upon BDNF stimulation. **a** Representative western blot analysis of FL-TrkB co-immunoprecipitation (IP) with SorCS2 from dorsal striatum of adult WT mice. Pulldown using normal IgG from the same species serves as a negative control. SorCS2 physically binds TrkB in dorsal striatum. **b** Quantification of ELISA signal from rat primary cultures (E18, DIV10) treated with 10 ng/ml BDNF for 15 min shows increased interaction between TrkB and SorCS2 upon BDNF stimulation. **c** Representative western blot analysis of total and phosphorylated TrkB from acute brain slices of dorsal striatum from P5 pups stimulated with 50 ng/ml BDNF for 1 hour. b-actin serves as a loading control. **d** Quantification of the immunoblots (**c**). TrkB phosphorylation in WT samples and SorCS2 KO samples revealed significant differences (One-way ANOVA, p=0.0008). When treated with BDNF, WT samples show an increase in pTrkB phosphorylation (p=0.0023), while SorCS2 KO mice exhibit decreased phosphorylation (p=0.0047) (Multiple comparison test with Tukey’s correction). The signal of pTrkB was normalized to total Full length TrkB. RBI = relative band intensity. Graphs show mean values ±SEM.

### SorCS2 promotes BDNF-dependent LTP in dorsal striatum

Given a critical role of BDNF-TrkB signaling in NMDAR-dependent LTP, we investigated if SorCS2 modifies synaptic potentiation in the corticostriatal synapses. We first examined α-amino-3-hydroxy-5-methyl-4-isoxazolepropionic acid receptor (AMPAR) activity by recording input-output (I/O) curves in the dorsal striatum in P35-45 mice. Field potentials were evoked in the white matter by stimulating the lower part of the corpus callosum directly overlying the dorsal striatum, and the synaptic response was quantified as the slope of the field excitatory postsynaptic potential (fEPSP). The I/O curves showed no differences in AMPAR activity between genotypes (**Figure 5a-b**). We then applied a theta burst stimulation (TBS) protocol for induction of LTP [40]. A stable baseline of at least 20 min preceded the TBS, and LTP was subsequently quantified as the percentage increase in the slope of the fEPSP compared to the baseline. Notably, whereas we observed robust LTP in WT slices for up to 180 min post stimulation (p<0.0001), LTP was substantially impaired in both the SorCS2 KO and R6/1 mice (p<0.0001 for both conditions) (**Figure 5c**). To explore whether the perturbed BDNF-TrkB signaling accounted for the blunted LTP in SorCS2 KO and R6/1 mice, we performed rescue experiments in which 10ng/ml BDNF was supplemented to the aCSF (**Figure 5d-g**). In WT mice, LTP was maintained although with reduced amplitude (**Figure 5e**). This is in accordance with studies reporting that acute exposure to BDNF increases basal synaptic potentiation and reduces LTP in WT mice [51]. Remarkably, BDNF fully restored LTP in SorCS2 KO slices as the synaptic strength was established at the same level as observed for untreated WT sections (**Figure 5f**). Hence, the attenuated BDNF-TrkB signaling in *Sorcs2-/-* mice can be overcome by addition of exogenous BDNF. On the contrary, BDNF failed to rescue the LTP in R6/1 mice. This data suggest that the functional deficits in the R6/1 mice are either a consequence of complete block in TrkB signaling or defects in BDNF-independent pathways (**Figure 5g**). All the measurements and statistics are summarized in (**Figure 5h**).

**Figure 5:**
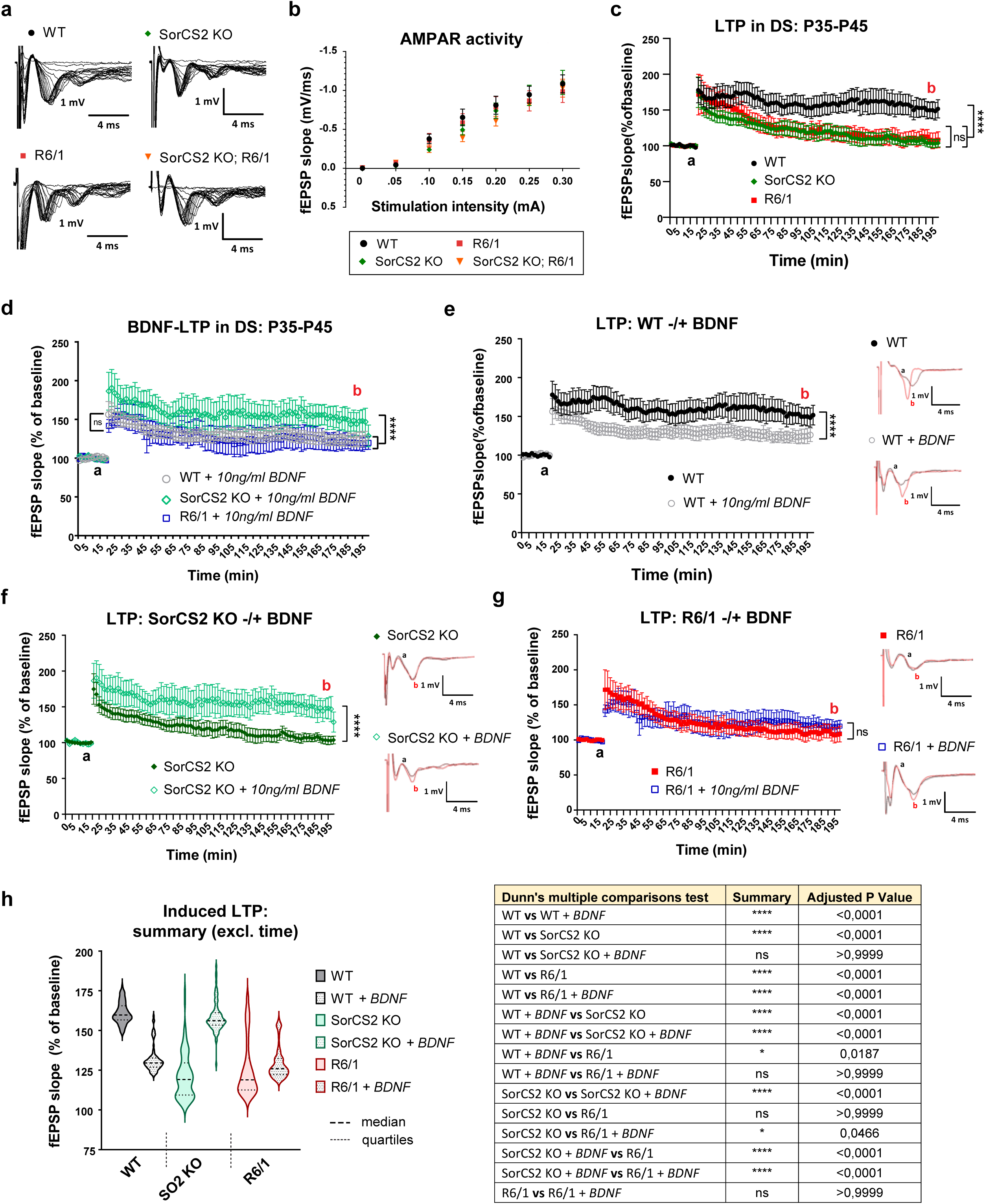
SorCS2 regulates BDNF-dependent LTP in dorsal striatum. **a** Representative traces of input/output curves for each genotyped that were measured in dorsal striatum of P35-45 males. **b** Corticostriatal synapses of p35-45 males show no significant differences in AMPAR activity. Unpaired two-tailed t-test. WT: n=4 (16 slices), SorCS2 KO: n=4 (16 slices), R6/1: n=4 (14 slices), SorCS2 KO; R6/1: n=4 (12 slices). **c** Induced LTP measurements in dorsal striatum of P35-45 males without treatment revealed impairments in SorCS2 KO and R6/1 models. Two-way ANOVA shows significant contribution of genotype and time (both p<0.0001). Kruskal-Wallis test and Dunn’s multiple comparisons results: SorCS2 KO vs WT (p<0.0001), R6/1 vs WT (p<0.0001). WT: n=9, SorCS2 KO: n=9, R6/1: n=7. **d** Induced LTP measurements in dorsal striatum of P35-45 males treated with 10ng/ml BDNF. BDNF stimulation rescued the deficient LTP in SorCS2 KO but not in R6/1. We observe a decrease in WT due to oversaturation of the LTP induction during the experiment. Two-way ANOVA revealed significant contribution of genotype and time (both p<0.0001). Kruskal-Wallis test and Dunn’s multiple comparisons results: SorCS2 KO vs WT (p<0.0001), R6/1 vs SorCS2 KO (p<0.0001). WT: n=8, SorCS2 KO: n=5, R6/1: n=6. **e-g** Comparison of the single genotypes –/+ BDNF. Two-way ANOVA revealed significant contribution of the genotype in (**e**) and (**f**) with p<0.0001, whereas R6/1 line shows no changes. **h** The violin plot visualizes the data for all conditions excluding the time factor and the baseline values. The data show significance p<0.0001 in Kruskal-Wallis test. Multiple-comparison testing is summarized in the table. **b-g** graphs show mean values ±SEM, **h** graph shows median with quartiles.

### Synaptic function of GluN2B is controlled by SorCS2

Synaptic accumulation and conductance of NMDA receptors is required for the induction of BDNF-dependent LTP [52]. We therefore compared NMDAR currents in the dorsal striatum of 35-45 days old SorCS2 KO and R6/1 mice. The recordings were performed in 0.1 mM magnesium to partially relieve the magnesium block of the NMDARs. Once AMPAR-and NMDAR-mediated potentials were stable, we first blocked AMPARs by adding the antagonist, cyanquixaline (DNQX) (**Figure 6a**). NMDAR potentials were then measured as the amplitude of the slow potential and normalized to the slope of the AMPAR-mediated potentials. We found that synaptic NMDAR-mediated currents were substantially attenuated in both SorCS2 KO and R6/1 mice (corresponding to approximately 60%), followed by a severe worsening of the phenotype (down to 31%) when the lines were crossbred (**Figure 6b**). Next, we explored the relative contribution of GluN2A and GluN2B to the total NMDAR conductivity. To this end, neurotransmission was recorded in brain slices treated with the selective GluN2B inhibitor RO 25-6981 maleate, followed by the GluN2A inhibitor PEAQX, respectively (**Figure 6a**). The GluN2B and GluN2A components were then calculated and normalized to the AMPAR currents for each experiment. Strikingly, we found that the attenuated NMDA currents in *Sorcs2^−/−^* animals were entirely accounted for by elimination of GluN2B neurotransmission (**Figure 6d**) with no impact on the GluN2A activity (**Figure 6c**). We measured a similar selective regulation in GluN2B activity in hippocampal CA3-CA1 synapses (**Suppl. Figure S5**), suggesting a more general function of SorCS2 in controlling NMDAR dependent neurotransmission. In marked contrast to SorCS2 KO, only GluN2A currents were affected in the dorsal striatum of the R6/1 mice (**Figure 6c-d**). These findings show that SorCS2 deficiency and the accumulation of mHTT affect the synaptic plasticity by different modes of action, which may explain why mice harboring both impairments have a more severe disease course.

**Figure 6:**
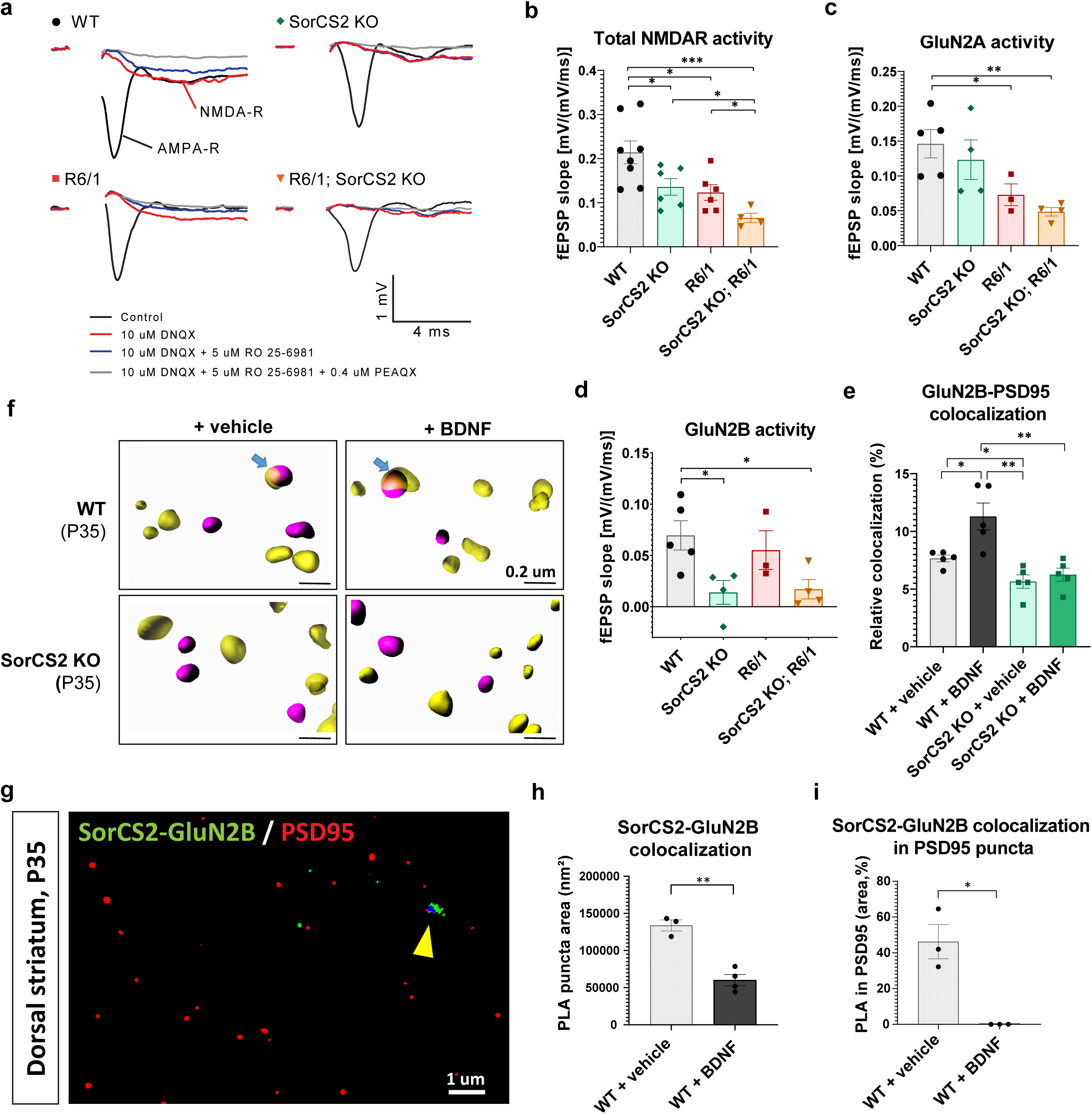
HD aggravation depends on SorCS2 targeting of GluN2B at the postsynaptic densities prior BDNF stimulation. **a** Field recordings of NMDA-R activity from *ex vivo* slices of dorsal striatum in P35-P45 transgenic mice. DNQX is AMPAR inhibitor, RO 25-6981 is GluN2B inhibitor, and PEAQX is GluN2A inhibitor were added sequentially. Brown-Forsythe ANOVA test and multiple-comparison test (MC) with Welch’s correction were used in (**b-e**). **b** Quantification of total NMDAR activity based on the field recordings show increased deficiency of NMDAR activity in the double-transgenic mice. ANOVA: p=0.0006; MC: SorCS2 KO vs WT (p=0.0309), R6/1 vs WT (p=0.0135), SorCS2 KO; R6/1 vs WT (p=0.0005), SorCS2 KO; R6/1 vs SorCS2 KO (p=0.0127), SorCS2 KO; R6/1 vs R6/1 (p=0.0243). **c** Quantification of GluN2A activity shows that the NMDAR deficiency is not dependent on GluN2A. ANOVA: p=0.0236. MC: R6/1 vs WT (p=0.0288), SorCS2 KO; R6/1 vs WT (p=0.0068). **d** Quantification of GluN2B activity revealed that deficient NMDAR activity is dependent of SorCS2-GluN2B functional interaction. ANOVA: p=0.0379; MC: SorCS2 KO vs WT (p=0.0191), SorCS2 KO; R6/1 vs WT (p=0.0194). **e** Quantification of super-resolution microscopy of GluN2B-PSD95 co-localization in *ex vivo* slices of dorsal striatum in presence or absence of BDNF. The results show that SorCS2 KO MSNs are deficient for GluN2B at the synapse, and they do not respond to BDNF stimulation. ANOVA: p=0.002, MC: WT-BDNF vs WT-vehicle (p=0.0339), SorCS2 KO-vehicle vs WT-vehicle (p=0.0214), SorCS2 KO-vehicle vs WT-BDNF (p=0.0053), SorCS2 KO-BDNF vs WT-BDNF (p=0.0088). **f** 3D reconstruction of the GluN2B (purple) co-localization (arrow) with PSD95 (yellow) at postsynaptic densities. The scale bar is 0.2 um. **g** Super-resolution Airyscan image of proximity ligation assay (PLA) for SorCS2-GluN2B interaction (in green) combined with immunofluorescence for PSD95 (in red) in *ex vivo* slices of dorsal striatum. Co-localization of SorCS2-GluN2B PLA signal with PSD95-positive objects are in blue (arrowhead). The scale bar is 1 um. **h** Quantification of SorCS2-GluN2B PLA signal in *ex vivo* slices of dorsal striatum treated with BDNF. There are fewer SorCS2-GluN2B protein complexes after BDNF stimulation. Welch’s t-test: p=0.0011. **i** Quantification of the presence of SorCS2-GluN2B complexes in PSD95-positive postsynaptic densities. SorCS2-GluN2B complexes are abolished at postsynaptic density after BDNF stimulation. Welch’s t-test: p=0.043.

### SorCS2 targets GluN2B to postsynaptic densities in a BDNF-dependent manner

To explore whether the effect of SorCS2 on synaptic plasticity was a consequence of altered expression of GluN2B or other key synaptic proteins, we performed western blotting of homogenates and synaptosomes from the dorsal striatum of WT and SorCS2 KO mice, age P35 (**Suppl. Figure S6a**). In the synaptosome fractions, we found no alterations in the NMDAR subunits GluN1, GluN2A, and GluN2B, neither in synaptophysin and pAkt levels (**Suppl. Figure S6b-d, g, k**). However, PSD95, TrkB, total Erk1/2, and pErk1/2 (but not the ratio between pErk and Erk1/2) were all increased in the homogenates (**Suppl. Figure S6e, f, h-j**), possibly to compensate for the reduced ability of BDNF to activate TrkB (*cf.* **Figure 4c-d**). Total Erk1/2 was increased in synaptosomes, but this did not translate into any alterations in Erk1/2 phosphorylation (**Suppl. Figure S6h-j**). Hence, the impaired GluN2B activity of corticostriatal synapses in the SorCS2 KO mice is not accounted for by altered expression of NMDARs or the selected synaptic proteins involved in the neurotransmission.

BDNF regulates the synaptic targeting and conductance of GluN2B in the corticostriatal pathway [18, 19, 53–55]. We therefore studied GluN2B expression in synapses of the dorsal striatum from SorCS2 KO mice with or without prior BDNF stimulation. The sections were immunostained for PSD95 and GluN2B, processed by Airy-scan super-resolution microscopy. Strikingly, GluN2B co-localization with PSD95 was lower by approximately 25% in the unstimulated SorCS2 KO slices compared to WT mice (**Figure 6e-f**). Furthermore, BDNF treatment increased postsynaptic enrichment of GluN2B by approximately 50% in WT mice (from 7.5% to 11.25%) but SorCS2 KO mice were completely refractory. Hence, it is likely that local activation of GluN2B already present at the synapse, rather than its recycling and recruitment from extrasynaptic sites, is accountable for the restoration of LTP by BDNF in SorCS2 deficient mice. These data suggest that SorCS2, aside from its ability to support TrkB signaling, also directly enables the synaptic targeting of GluN2B.

Finally, we asked whether SorCS2, similarly to its interaction with TrkB, might physically interact with GluN2B in a BDNF-dependent manner. To this end, we used proximity ligation assay (PLA) for SorCS2 and GluN2B interaction. This method labels proteins that are closely juxtapositioned. Immunofluorescence microscopy revealed that the receptors co-localize throughout MSNs in naïve WT slices. Strikingly, stimulation with BDNF reduced the SorCS2-GluN2B interaction in MSNs by 60% indicating disengagement of the SorCS2-GluN2B complex (**Figure 6g-h**). Remarkably, almost 50% of the SorCS2-GluN2B PLA puncta were present in PSD95-positive domains in the unstimulated dorsal striatum, but BDNF treatment completely abolished the synaptic co-localization (**Figure 6i**). Taken together, the data demonstrate that SorCS2 binds GluN2B to enable its translocation to the synapse, after which BDNF abolishes their heterodimerization. This may stabilize the synaptic expression of the NMDAR subunit and condition it for neurotransmssion.

## DISCUSSION

In the healthy brain, BDNF-TrkB signaling serves two functions; it enables the activity-dependent regulation of synaptic structures and function, and it supports the neuronal integrity through trophic stimulation. A cornerstone in the HD neuropathology is the diminished BDNF activity as consequence of reduced BDNF production or its impaired anterograde transport from cortex to striatum, or decreased expression or signalling abilities of TrkB. At early stages of the disease, altered NMDA receptor activity caused by reduced BDNF-TrkB signaling is considered fundamental while at later stages lowered neuroprotection may propel the pathophysiological changes [56, 57]. Accordingly, some HD mouse models develop overt neurodegeneration, whereas in other models synaptic disturbances such as disrupted NMDA-dependent LTP manifest but without apparent loss of neurons [8, 28, 58]. Here we identified SorC2 as a molecular hub that regulates BDNF signaling, synaptic plasticity, and HD progression through dynamic changes in its physical interactions with TrkB and GluN2B.

We show that SorCS2 is selectively expressed by MSNs in dorsal striatum, and that the performance of SorCS2 KO mice on the rotarod is compromised. Most likely, this effect was not accounted for by neuronal cell loss since the surrogate parameters of neuronal degeneration DARPP-32 and clasping were not altered in the aged SorCS2 knockouts. On the other hand, SorCS2 deficiency on the R6/1 genetic background severely aggravated the motor phenotypes and accelerated the disease progression. One of the earliest alterations during HD development is a dysfunction of corticostriatal synapses [56]. In HD mouse models such as BACHD, Q175, or HdhQ7/Q111, the motor deficits have been attributed to the loss of LTP [8, 59]. BDNF signalling is fundamental for ionotropic potentiation, and both GluN2A and GluN2B are required for LTP in corticostriatal synapses [8, 60, 61]. In hippocampal neurons, the activation of TrkB results in phosphorylation of GluN1 [62] and GluN2B subunits [18, 19], but not GluN2A [63], which is required for synaptic trafficking of NMDA receptors and LTP induction [53]. We found that SorCS2 binds TrkB in BDNF-dependent manner, and that this interaction is required for efficient activation of TrkB. Furthermore, the absence of SorCS2 caused a significant decrease in TrkB phosphorylation and abolished LTP in the corticostriatal pathway. These features were probably caused by a severe deficiency in GluN2B currents as the neurotransmission by GluN2A was preserved. Our findings contrast those of *Ma et al 2017* who reported that GluN2A expression is decreased in MSNs from SorCS2 KO mice. However, the authors did not study any functional consequences of their observation [64]. Although we cannot discount the possibility that SorCS2 affects the subcellular distribution of GluN2A, it does not seem to be of major functional significance. In contrast to SorCS2 deficient mice, GluN2A currents were affected only in the R6/1 line. This is in accordance with a recent study that reported reduced LTP, impaired GluN2A but unaltered GluN2B activity in the hippocampus of R6/1 mice [65]. In our SorCS2 KO; R6/1 double transgenic mice, GluN2A and GluN2B were both severely attenuated which translated into a marked reduction in total NMDA receptor currents. This observation may explain the worsening of the behavioral phenotypes in our double transgenic mice.

Aside from the function of SorCS2 in TrkB activation, which induces phosphorylation of GluN2B and GluN1 that is necessary for their synaptic enrichment, SorCS2 also modulates GluN2B function through a direct, BDNF-independent interaction. Co-immunoprecipitation experiments recently showed that SorCS2 and GluN2B form heterodimers in hippocampal extracts [66]. We found that synaptic clustering of GluN2B in the dorsal striatum from SorCS2 KO was reduced by 25% compared to WT mice. Furthermore, GluN2B currents in naïve, unstimulated slices were diminished by 80% in SorCS2 KO while GluN2A currents remained unaffected. Strikingly, the interaction between SorCS2 and GluN2B was dynamic in MSNs as BDNF disrupted the complexes. SorCS2 is engaged in several aspects of protein sorting directing its cargo between Golgi, cell surface and endosomes [36]. Notably we previously found that SorCS2 facilitates the synaptic translocation of TrkB to postsynaptic densities in hippocampal neurons [33]. Given its many trafficking pathways, SorCS2 could potentially enable the synaptic insertion of GluN2B during exocytosis, its lateral diffusion between the synapse and extrasynaptic sites, or its endocytosis and recycling. Even though the sorting pathway is currently unclear, our study identifies SorCS2 as a novel sorting mechanism to regulate the postsynaptic membrane localization of GluN2B.

Psychiatric symptoms are common in HD [1]. Our previous studies discovered that SorCS2 KO mice exhibit cognitive impairments and traits typifying ADHD and schizophrenia [33, 37]. Similarly to the dorsal striatum, ionotropic LTP was absent in CA3-CA1 synapses but could be rescued by exogenous BDNF. Here we find that GluN2B but not GluN2A currents are substantially reduced also in CA3-CA1 synapses, suggesting a broader function of SorCS2 in modifying NMDAR-dependent plasticity. SorCS2 plays an important role for dopaminergic functionality as SorCS2 deficiency severely perturbs dopaminergic firing rates in the ventral tegmental area, striatal dopamine metabolism, and dopamine D1 receptor (D1R) sensitivity [35, 37]. Inquiringly, D1R cooperates with TrkB to modulate ERK signalling in striatal neurons and to increase GluN2B phosphorylation [67, 68]. Given that dysfunctional striatal dopamine signalling contributes to the HD pathophysiology in mouse models [69], it is possible that yet another loop may be added to the many functions of SorCS2 in Huntington’s disease.

SorCS2 mRNA and protein levels were reduced in the dorsal striatum of symptomatic R6/1 mice with no changes in cortex. Interestingly, the expression level was dependent on age and HD progression as it was not detected in young asymptomatic mice. A separate study reported reduced SorCS2 expression also in zQ17Z and R6/2 HD mice [64]. Inversely, others found upregulation of SorCS2 mRNA in MSNs in response to deep-brain stimulation in a Parkinson’s disease mouse model [70]. It appears that SorCS2 expression in MSNs is highly controlled by internal and external stimuli, and can dynamically change, e.g. as a reaction to pathophysiological changes in the tissue. A recent study reported that mHTT can bind SorCS2 leading to its redistribution and aggregation in MSNs [64]. Hence, in addition to attenuated gene transcription, scavenging of SorCS2 protein by mHTT may further reduce its expression in MSNs.

Taken together, we provide evidence that SorCS2 operates by two mechanisms to orchestrate GluN2B-dependent plasticity in MSNs. First, it associates with TrkB to enable its activation by BDNF, which is required to phosphorylate GluN2B and to empower GluN2B enrichment at the postsynaptic densities. However, in parallel to this, SorCS2 may assist in targeting GluN2B to the synapse through their direct interaction. Once at the synapse, BDNF uncouples SorCS2 from GluN2B, which may condition GluN2B for the required biological activity. The dynamic interaction of SorCS2 with TrkB, and GluN2B, respectively, might thus produce an amplifying loop where SorCS2, which is liberated from GluN2B by BDNF, can be reused by TrkB to further fuel BDNF signaling, phosphorylation of GluN2B, and the induction of LTP. This proposed model is schematized in the **Figure 7**. Even though the primary determinant of HD onset is the length of the CAG repeats on the expanded allele [1, 71], yet this only accounts for 50-70% variations in the age of onset [30, 31, 71, 72] . Notably, a recent GWAS study identified SNPs and missense mutations in *SORCS2* associated with HD [32], and another study has reported reduced levels of SorCS2 mRNA in post-mortem brains of HD patients [73]. Here we have provided mechanistic support for *SORCS2* as one of the long-sought disease modifying genes in Huntington’s disease.

**Figure 7:**
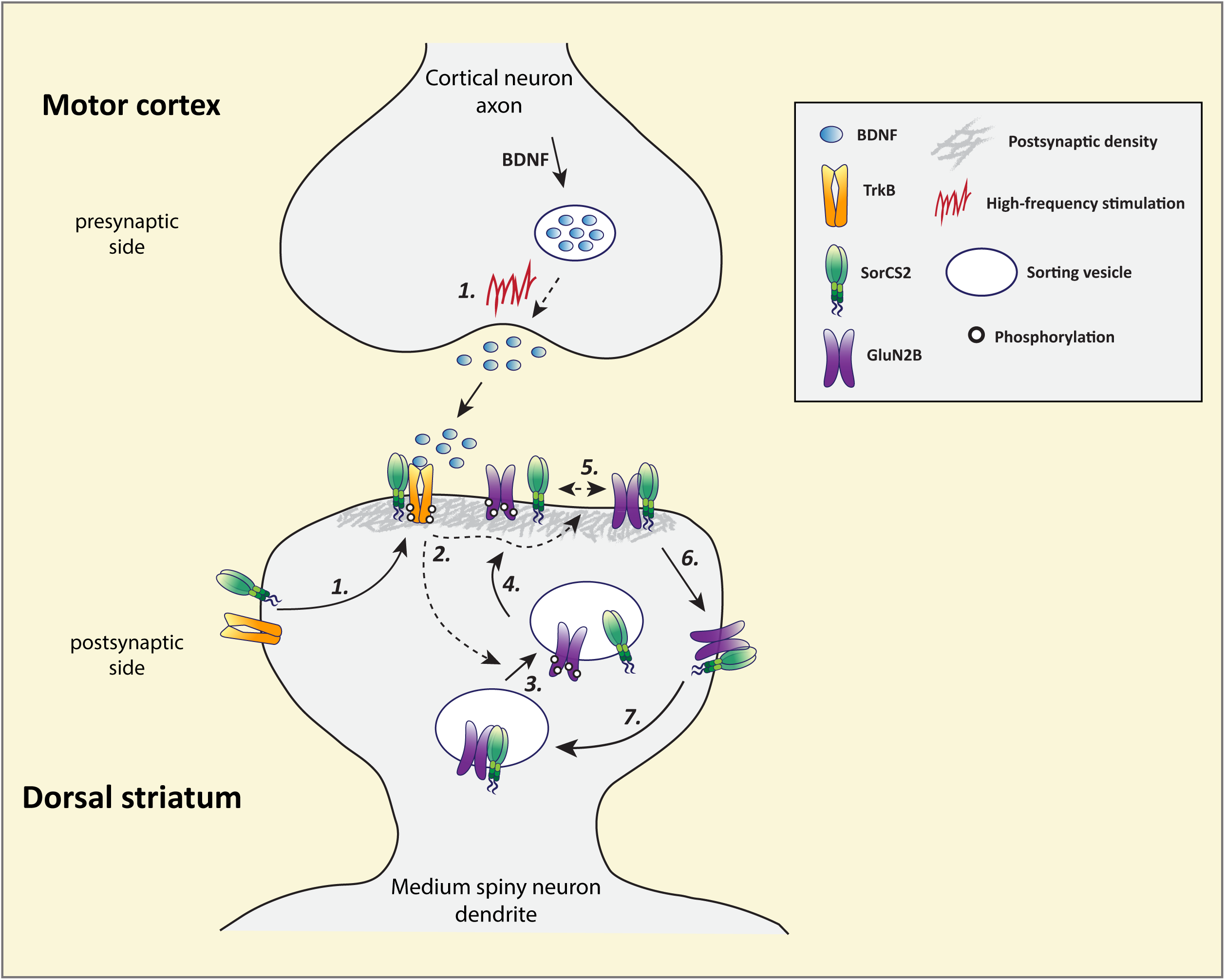
A schematic model depicting the role of SorCS2 in neurotransmission of corticostriatal synapses. **Step 1.** Upon BDNF release from cortical afferents, SorCS2 binds TrkB upon which they translocate to postsynaptic densities [33]. TrkB dimerization is strengthened by BDNF which is required for the robust phosphorylation of TrkB. **Step 2-3.** Activation of TrkB leads to dissociation of SorCS2 from GluN2B, possibly as consequence of GluN2B phosphorylation. **Step 4.** Once liberated from SorCS2, GluN2B is targeted to postsynaptic densities directly (displayed here) or after being in transit in the perisynapse (not displayed). **Step 5.** When GluN2B phosphorylation is not being maintained by BDNF, GluN2B will physically interact with SorCS2 in neuronal structures including postsynaptic densities. **Step 6-7.** SorCS2-GluN2B complexes may potentially be endocytosed into recycling vesicles following the lateral diffusion to the perisynapse.

## CONCLUSIONS

Reduced BDNF-TrkB signaling and NMDAR-dependent synaptic plasticity are hallmarks of early stages of Huntington’s disease. We show that SorCS2 enables TrkB activation and navigates GluN2B to MSN synapses. The binding between SorCS2 and TrkB and GluN2B, respectively are mutually exclusive and controlled by BDNF. While BDNF stimulates complex formation between SorCS2 and TrkB to amplify BDNF signaling, it uncouples SorCS2 from GluN2B resulting in enrichment of the neurotransmitter subunit at postsynaptic densities. During HD progression SorCS2 expression declines which compromises BDNF-TrkB signaling and NMDAR-dependent plasticity thereby accelerating the disease course. These findings may nominate SorCS2 as therapeutic target to slow down the progression of Huntingtońs disease.

## LIST OF ABBREVIATIONS

aCSF: artificial cerebral spinal fluid
AMPAR: α-amino-3-hydroxy-5-methyl-4-isoxazolepropionic acid receptor
BDNF: Brain-derived neurotrophic factor
Cf.: cross reference
D1R: dopamine D1 receptor
DS: dorsal striatum
fEPSP: field excitatory postsynaptic potential
GFAP: Glial fibrillary acidic protein
GluN1/2B/2A: Glutamate Ionotropic Receptor NMDA Type Subunit 1/2B/2A
GWAS: genome-wide association study
HD: Huntington’s disease
HTT/mHTT: Huntingtin / mutated Huntingtin
I/O: input-output curves
IHC: in situ hybridization
IP: immunoprecipitation
KO: knock out
LTP: long-term potentiation
MC: multi comparison test
MSNs: medium spiny neurons
NMDAR: N-methyl-D-aspartate receptor
PLA: proximity ligation assay
proBDNF: precursor of Brain-derived neurotrophic factor
qPCR: real-time quantitative polymerase chain reaction
RBI: relative band intensity
SEM: standard error of mean
SNP: single nucleotide polymorphism
SorCS2: Sortilin Related VPS10 Domain Containing Receptor 2
TBS: theta burst stimulation
TrkB: Tropomyosin related kinase receptor B
TrkB-FL: full length Tropomyosin related kinase receptor B
TrkB-T.1: Truncated variant of Tropomyosin related kinase receptor B
WT: wild type

## DECLARATIONS

### Ethics approval

Animal experiments were performed in full compliance with Danish and European regulations. All experiments are covered by permissions: 2012-15-2934-00397 (C3) and 2017-15-0201-01192 (C3) issued by the Danish Animal Experiments Inspectorate.

### Consent for publication

Not applicable

### Availability of data and materials

Not applicable

### Authors’ contributions

AS performed some experiments, analyzed the data, wrote the manuscript, prepared all the figures, and coordinated finalization of the study; this determined her to become the first shared first-author. NSD planned and conducted some experiments, analyzed the data, and wrote a manuscript draft. MP performed SR microscopy and analyzed the data. MP provided unique insight into the molecular mechanism, and this is why he is the first shared-second author. NKM helped with behavior experiments, built the catwalk assay, and developed MATLAB programs for analyzing behavioral data. PLO helped with synaptosomes preparation and electrophysiological measurements. BV helped with conducting the mice work and performed some biochemical experiments. SLB and JCA measured BDNF and proBDNF levels. PC and EC provided the ELISA data for TrkB-SorCS2 interaction. AU analyzed the synaptic proteins in synaptosomal extraction. LK isolated the synaptosomes and performed some pioneer pulldown experiments. LW performed qPCR. IF and SN helped with *in vivo* experiments that were not included in the final study. MMH provided input and support to the electrophysiological experiments. MFK trained NSD in performing biochemical experiments and drafting the manuscript. UB trained NSD, and performed some electrophysiological experiments. AN designed the study, coordinated the research, and wrote the manuscript.

## Supporting information

Supplementary video SV1 - WT mice

Supplementary video SV4 - gait of SorCS2 KO; R6/1 mice

Supplementary video SV3 - gait of R6/1 mice

Supplementary video SV2 - gait of SorCS2 KO mice

## Acknowledgements

We acknowledge Kristian Skipper Alsberg and Anne Kathrine Aalling Sørensen for their technical involvement in part of the project, which was not included in this manuscript. We thank Sune Skeldal and Anne Kerstine Glintborg Jensen for the technical help, and Stella Solveig Nolte for coordinating the mice work, and for the genotyping. We thank the department of Biomedicine at Aarhus University for providing the infrastructure and administrative support.

## ADDITIONAL MATERIALS

Supplementary video SV1: Representative video of gait assay result for WT mice walking through the catwalk assay.

Supplementary video SV2: Representative video of SorCS2 KO mouse line walking through the catwalk assay.

Supplementary video SV3: Representative video of R6/1 mouse line walking through the catwalk assay.

Supplementary video SV4: Representative video of SorCS2 KO; R6/1 mouse line walking through the catwalk assay.

---

**Supplementary Figure S1:**
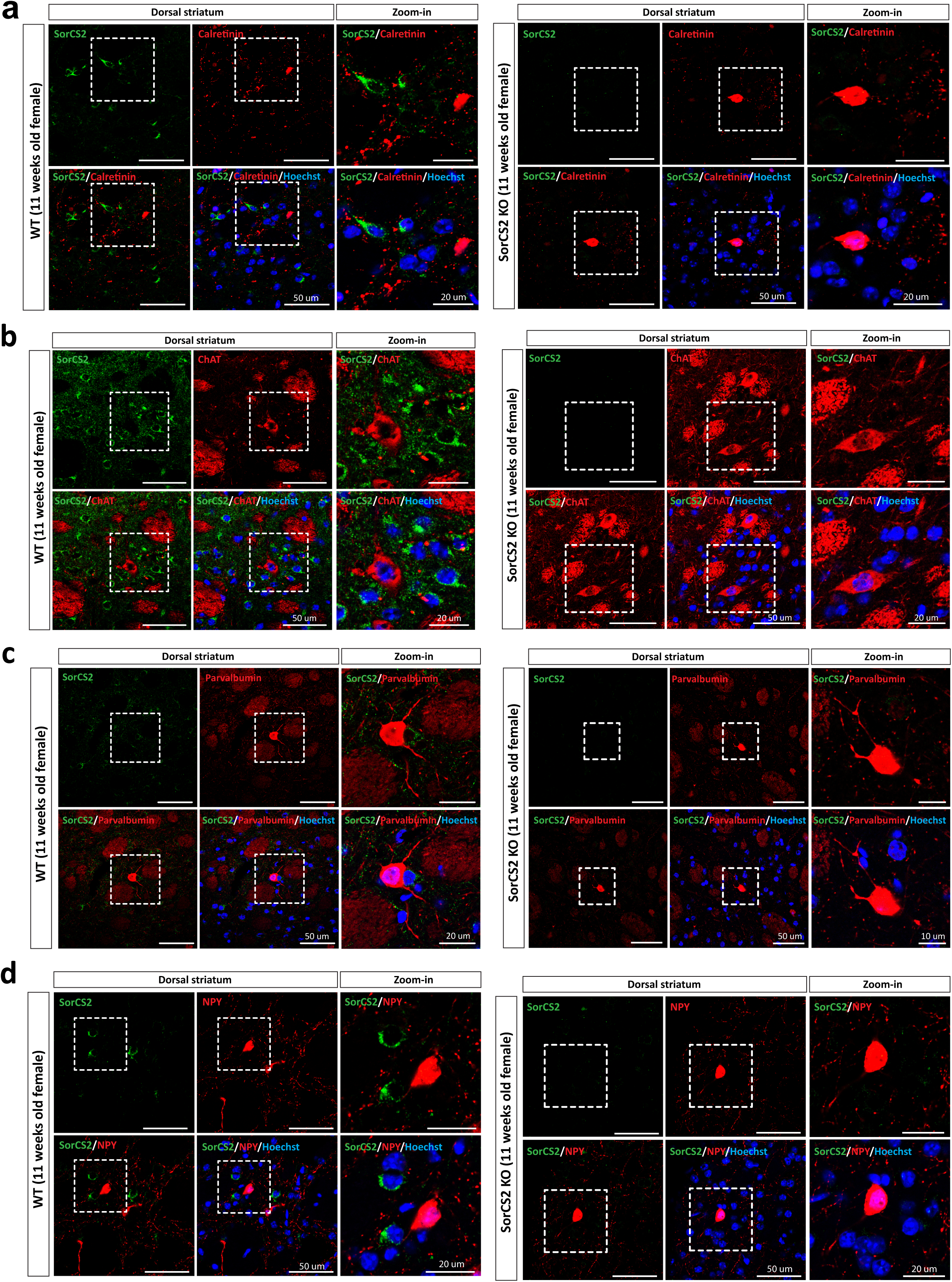
SorCS2 is not expressed by interneurons in dorsal striatum. Immunofluorescence labelling four major interneuron populations in dorsal striatum followed by confocal microscopy. SorCS2 KO served as a negative control to which we calibrated the imaging settings. We used adult females for this analysis. **a** SorCS2 (in green) is not co-expressed in Calretinin (in red) interneurons. **b** SorCS2 is not expressed by large-aspiny cholinergic interneurons expressing ChaT. **c** SorCS2 is not expressed by GABAerfic fast-spiking parvalbumine-positive interneurons. **d** SorCS2 is not expressed by low-threshold-spiking NPY positive interneurons. Hoechst staining (in blue) labels the cell nuclei. The scale bar corresponds to 50µm, and to 20µm in magnified figures.

**Supplementary Figure S2:**
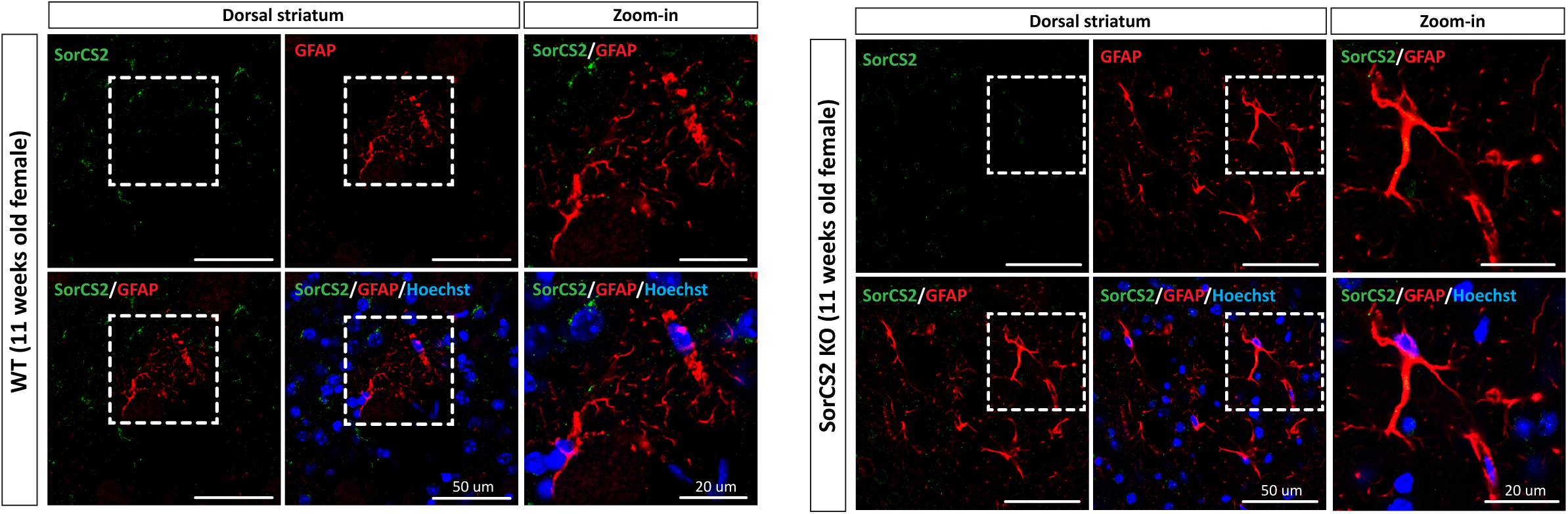
SorCS2 is not expressed by glial cells. Immunofluorescence labelling of SorCS2 (in green) and GFAP+ glial cells (in red) in dorsal striatum followed by confocal microscopy. SorCS2 KO served as a negative control to which we calibrated the imaging settings. We used adult females for this analysis. Hoechst staining (in blue) labels the cell nuclei. The scale bar corresponds to 50µm, and to 20µm in magnified figures.

**Supplementary Figure S3:**
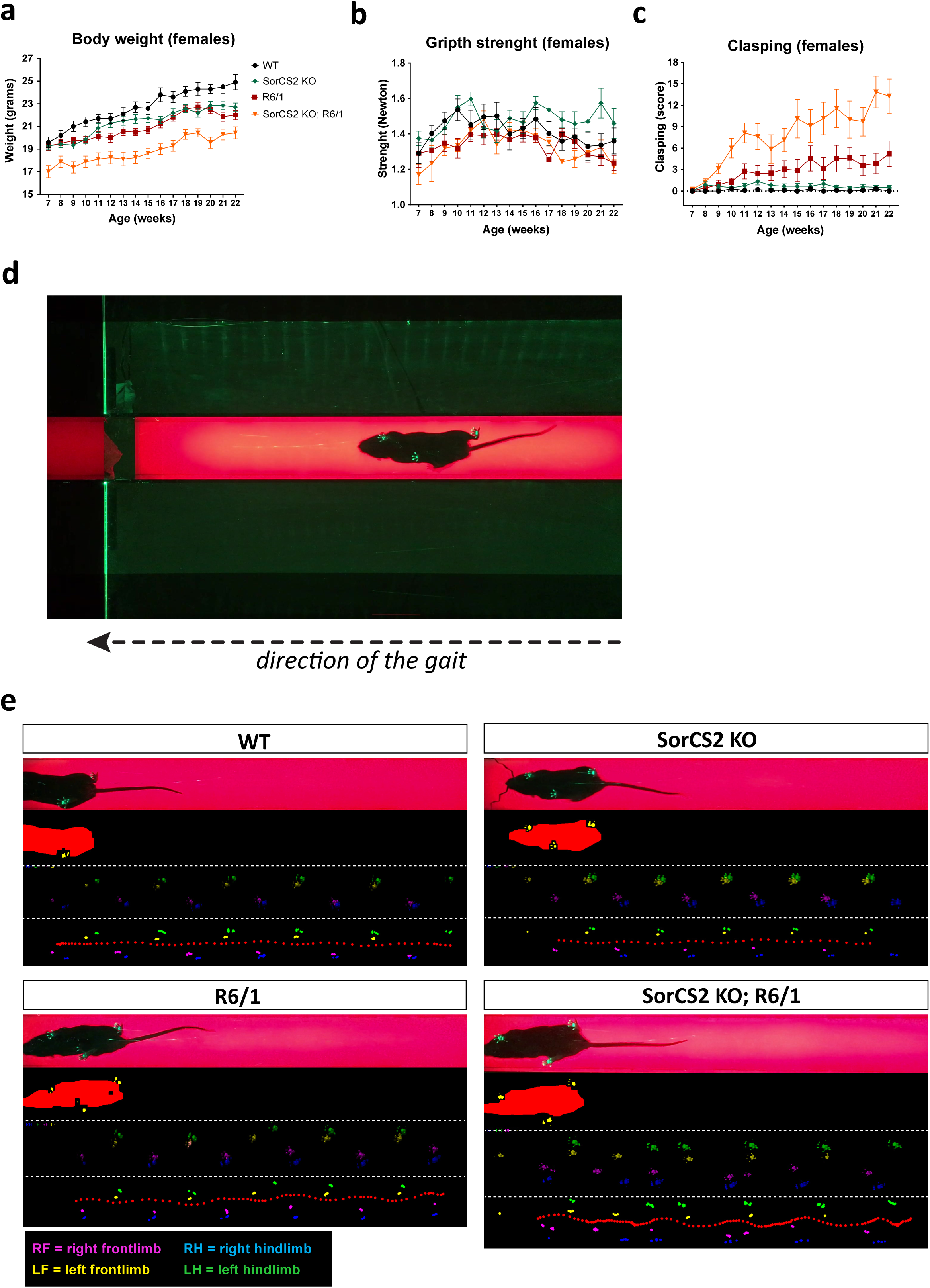
Physiological and behavior monitoring of transgenic HD mice including females. **a** Body weight in females with significant contribution of genotype and age (both p<0.0001, Mixed-effects model analysis with Tukey’s correction). Repeated measurements one-way ANOVA and Tukey’s multiple comparisons test: SorCS2 KO vs WT (p<0.0001), R6/1 vs WT (p<0.0001), SorCS2 KO; R6/1 vs WT (p<0.0001), R6/1 vs SorCS2 KO (p=0.0267), SorCS2 KO; R6/1 vs SorCS2 KO (p<0.0001), SorCS2 KO; R6/1 vs R6/1 (p<0.0001). **b** Grip strength in females with significant contribution of genotype (p<0.0001) and age (p=0.001) (Mixed-effects model analysis with Tukey’s correction). Repeated measurements one-way ANOVA and Tukey’s multiple comparisons test: SorCS2 KO vs WT (p=0.0441), R6/1 vs WT (p=0.0009), SorCS2 KO; R6/1 vs WT (p=0.0006), R6/1 vs SorCS2 KO (p<0.0001), SorCS2 KO; R6/1 vs SorCS2 KO (p<0.0001). **c** Clasping score in females with significant contribution of genotype and age (both p<0.0001, Mixed-effects model analysis with with Tukey’s correction). Repeated measurements one-way ANOVA and Tukey’s multiple comparisons test: SorCS2 KO vs WT (p<0.0001), R6/1 vs WT (p<0.0001), SorCS2 KO; R6/1 vs WT (p<0.0001), R6/1 vs SorCS2 KO (p=0.0003), SorCS2 KO; R6/1 vs SorCS2 KO (p<0.0001), SorCS2 KO; R6/1 vs R6/1 (p<0.0001). **a-c** WT: n=10, SorCS2 KO: n=14, R6/1: n=16, SorCS2 KO; R6/1: n=8. **D** A built-in-house catwalk apparatus for a gait assay with a mouse inside of the corridor walking towards the box with the bedding. The paws reflect the green light. **e** Representative images of the gait from the different genotypes with decoded paws and visualized gait pattern. R6/1 have impaired gait which worsens in SorCS2 KO; R6/1 mice. RBI = Relative band intensity.

**Supplementary Figure S4:**
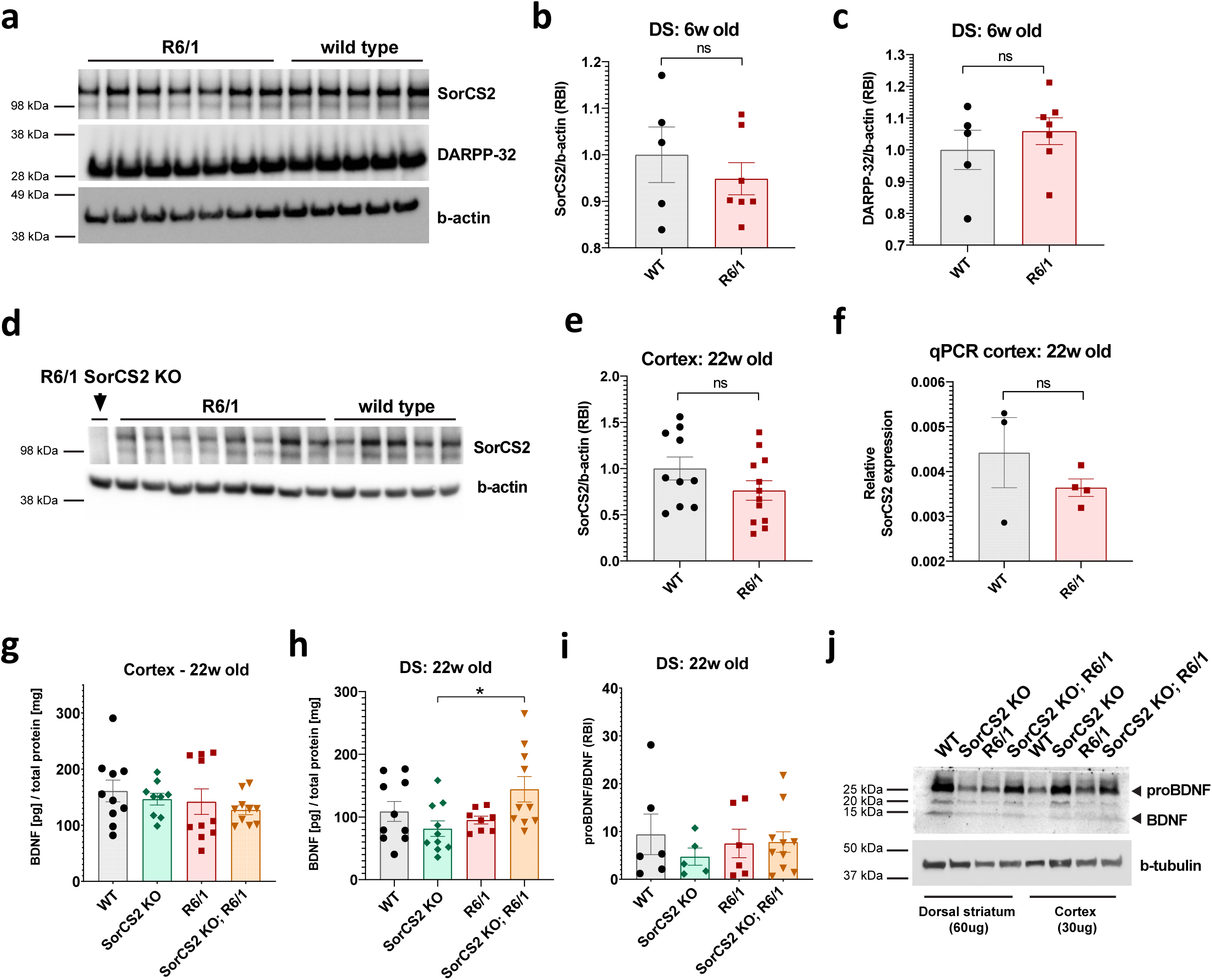
Protein analysis of cortex and dorsal striatum. **a** Representative immunoblot of SorCS2 and DARPP-32 in the dorsal striatum of P42-44 mice, which is quantified via densitometry of SorCS2 (**b**) and DARPP-32 (**c**) bands normalized to b-actin. We detected no differences in SorCS2 and DARPP-32 at this stage (unpaired two-tailed t-test). **d** Representative immunoblot showing SorCS2 protein levels in cortex of WT and R6/1 lines which do not differ quantified in **e** (unpaired two-tailed t-test). **f** Quantitative real-time PCR detects SorCS2 in cortex from 22-weeks-old mice. The expression is normalized to GAPDH transcripts serving as a housekeeping gene. There is no difference between WT and R6/1 line (unpaired two-tailed t-test). **g** BDNF levels in cortex from 22-week-old mice determined by ELISA. There is no significant differences between the genotypes (ordinary one-way ANOVA with Tukey’s multiple comparisons test). **h** BDNF levels in dorsal striatum from 22-week-old mice determined by ELISA. We detected upregulated levels of BDNF in the double-transgenic HD model. Significant change in ordinary one-way ANOVA (p=0.0357). Tukey’s multiple comparisons test: SorCS2 KO; R6/1 vs SorCS2 KO (p=0.0265). **i** Quantification of the ratio between proBDNF and BDNF levels was determined by densitometry of western blots in dorsal striatum from 22-week-old mice. There was no difference between the genotypes (ordinary one-way ANOVA with Tukey’s multiple comparisons test). **j** Representative immunoblot of proBDNF and BDNF in dorsal striatum in 22-week-old mice. All graphs show mean with ±SEM.

**Supplementary Figure S5:**
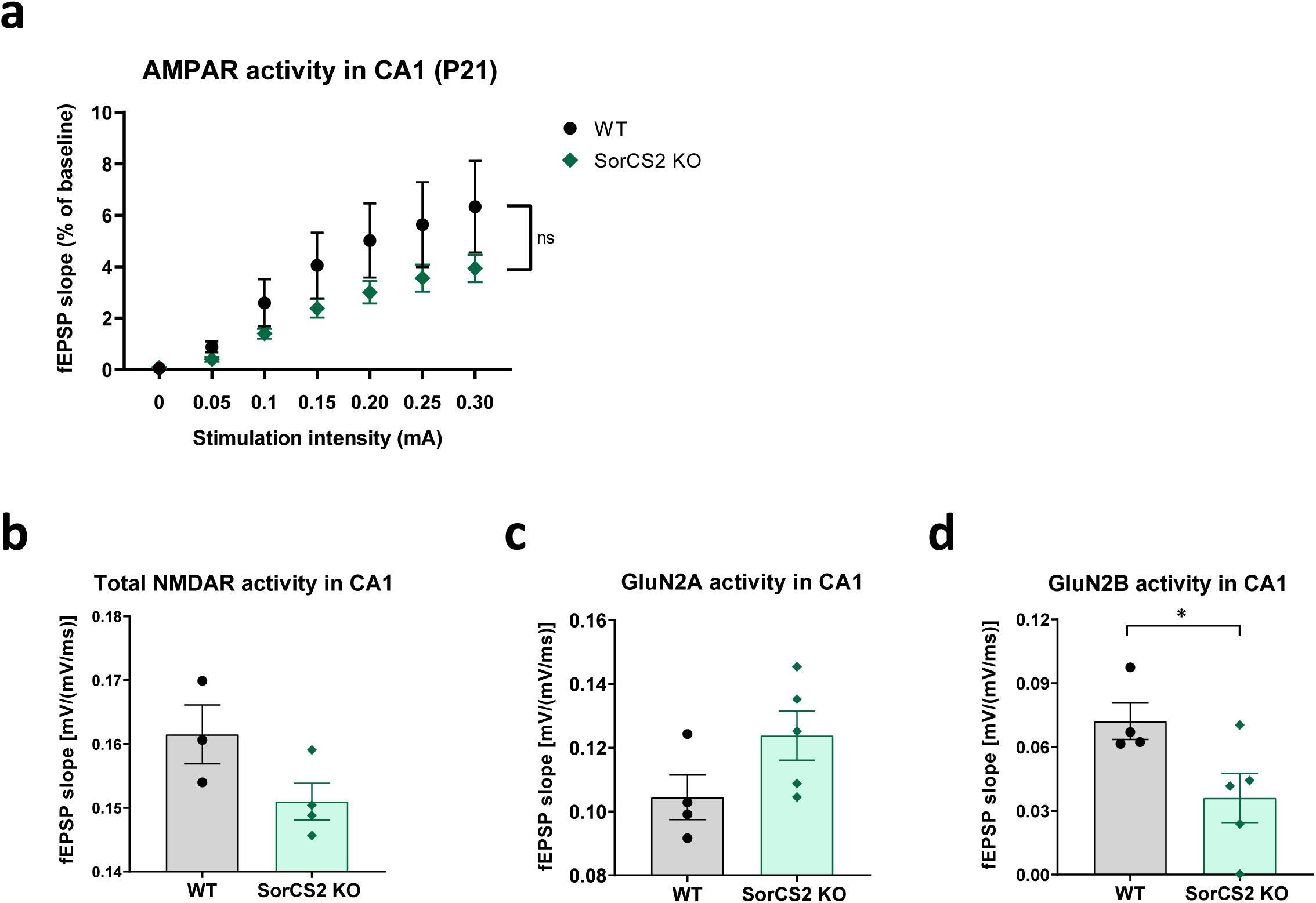
Analysis of AMPAR and NMDAR activity in CA3-CA1 synapses of hippocampus. **a** Recordings of field potentials from CA3-CA1 synapses show no significant difference in AMPAR activity in regards to genotype (two-way ANOVA, WT: n=5, SorCS2 KO: n=8). Quantification of total NMDAR activity (**b**) and GluN2A activity (**c**) revealed no difference; however, SorCS2 KO mice have decreased GluN2B activity (p=0.0419) as shown in (**d**). Unpaired Welch’s t-test. Mice were P21-22 of age. All graphs shows mean values ± SEM.

**Supplementary Figure S6:**
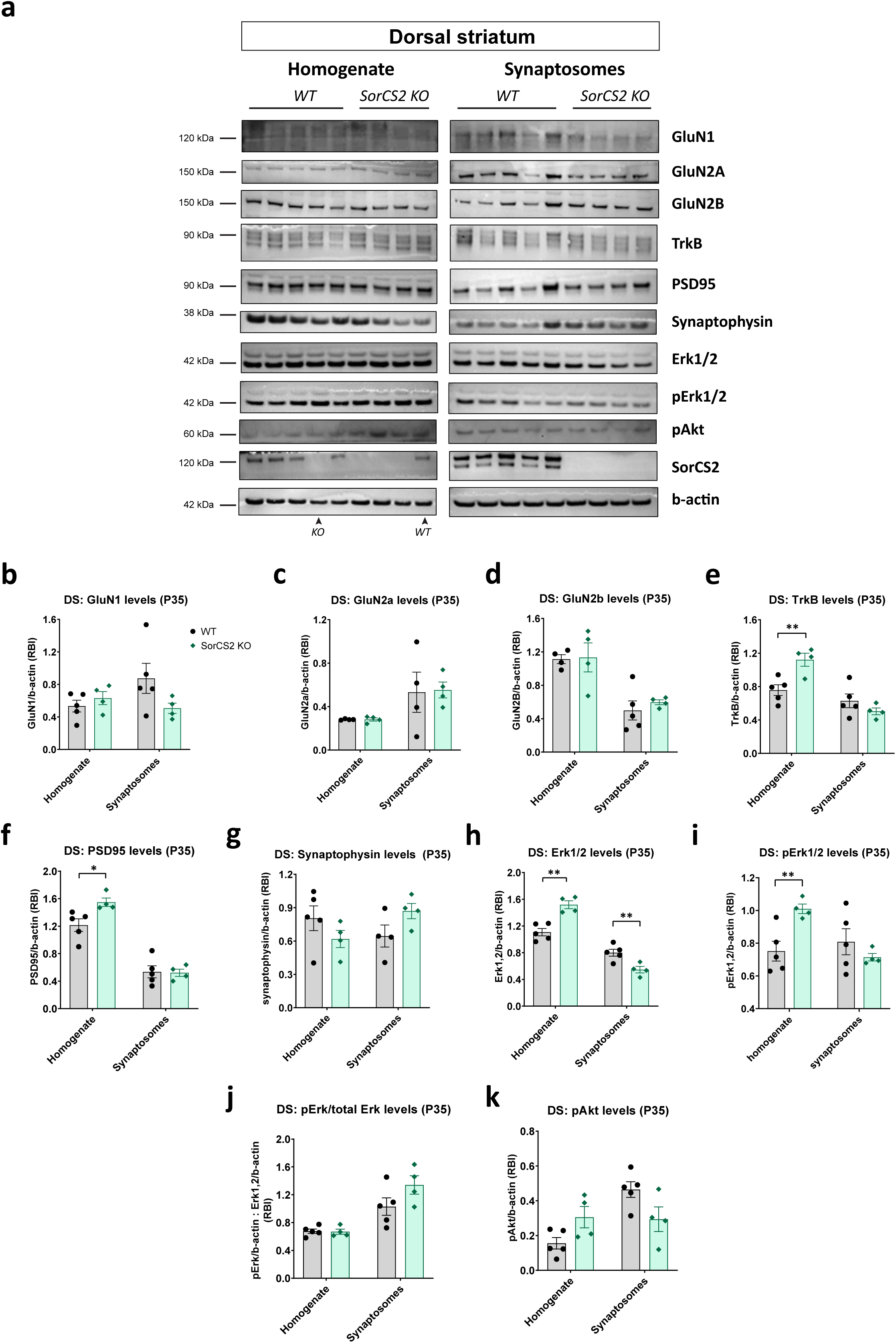
Analysis of selected synaptic proteins in homogenates and synaptosomes of dorsal striatum from P35 WT and SorCS2 KO mice. **a** Immunoblots of selected synaptic proteins in two distinct protein fractions. These data are quantified in the below graphs using densitometry and normalized to b-actin, followed by unpaired two-tailed t-test. We detect no difference in GluN1 (**b**), GluN2A (**c**) neither in GluN2B (**d**). As reported in Figure 3B, we detect upregulated levels of TrkB in the SorCS2 KO homogenate (**e**) (p=0.010742). We observe upregulation of PSD95 in SorCS2 KO homogenate (p=0.020426), however synaptosomes reflect normal levels of PSD95 (**f**). **g** Synaptophysin levels tend to decrease in SorCS2 KO homogenate and increase in synaptosomes, but there is no statistical significance. **h** We observe significant increase in total Erk1,2 in SorCS2 KO homogenate (p=0.0016), and a decrease in synaptosomes (p=0.008992). Mice were P35 of age. RBI = Relative band intensity. All graphs show mean ± SEM.

## REFERENCES

1. Ross CA, Tabrizi SJ: Huntington’s disease: from molecular pathogenesis to clinical treatment. The Lancet Neurology 2011, 10:83–98.

2. Jimenez-Sanchez M, Licitra F, Underwood BR, Rubinsztein DC: Huntington’s Disease: Mechanisms of Pathogenesis and Therapeutic Strategies. Cold Spring Harbor perspectives in medicine 2017, 7.

3. Bates GP, Dorsey R, Gusella JF, Hayden MR, Kay C, Leavitt BR, Nance M, Ross CA, Scahill RI, Wetzel R, et al: Huntington disease. Nature reviews Disease primers 2015, 1:15005.

4. McColgan P, Tabrizi SJ: Huntington’s disease: a clinical review. Eur J Neurol 2018, 25:24–34.

5. Zuccato C, Ciammola A, Rigamonti D, Leavitt BR, Goffredo D, Conti L, MacDonald ME, Friedlander RM, Silani V, Hayden MR, et al: Loss of huntingtin-mediated BDNF gene transcription in Huntington’s disease. Science (New York, NY) 2001, 293:493–498.

6. Gauthier LR, Charrin BC, Borrell-Pagès M, Dompierre JP, Rangone H, Cordelières FP, De Mey J, MacDonald ME, Lessmann V, Humbert S, Saudou F: Huntingtin controls neurotrophic support and survival of neurons by enhancing BDNF vesicular transport along microtubules. Cell 2004, 118:127–138.

7. Liot G, Zala D, Pla P, Mottet G, Piel M, Saudou F: Mutant Huntingtin alters retrograde transport of TrkB receptors in striatal dendrites. The Journal of neuroscience : the official journal of the Society for Neuroscience 2013, 33:6298–6309.

8. Plotkin JL, Day M, Peterson JD, Xie Z, Kress GJ, Rafalovich I, Kondapalli J, Gertler TS, Flajolet M, Greengard P, et al: Impaired TrkB receptor signaling underlies corticostriatal dysfunction in Huntington’s disease. Neuron 2014, 83:178–188.

9. Jiang M, Peng Q, Liu X, Jin J, Hou Z, Zhang J, Mori S, Ross CA, Ye K, Duan W: Small-molecule TrkB receptor agonists improve motor function and extend survival in a mouse model of Huntington’s disease. Hum Mol Genet 2013, 22:2462–2470.

10. Lee JM, Ramos EM, Lee JH, Gillis T, Mysore JS, Hayden MR, Warby SC, Morrison P, Nance M, Ross CA, et al: CAG repeat expansion in Huntington disease determines age at onset in a fully dominant fashion. Neurology 2012, 78:690–695.

11. Gusella JF, MacDonald ME, Lee JM: Genetic modifiers of Huntington’s disease. Movement disorders : official journal of the Movement Disorder Society 2014, 29:1359–1365.

12. Zuccato C, Cattaneo E: Role of brain-derived neurotrophic factor in Huntington’s disease. Prog Neurobiol 2007, 81:294–330.

13. Baydyuk M, Xu B: BDNF signaling and survival of striatal neurons. Frontiers in cellular neuroscience 2014, 8:254.

14. Minichiello L: TrkB signalling pathways in LTP and learning. Nature reviews Neuroscience 2009, 10:850–860.

15. Panja D, Bramham CR: BDNF mechanisms in late LTP formation: A synthesis and breakdown. Neuropharmacology 2014, 76 Pt C:664–676.

16. Leal G, Bramham CR, Duarte CB: BDNF and Hippocampal Synaptic Plasticity. Vitam Horm 2017, 104:153–195.

17. Carvalho AL, Caldeira MV, Santos SD, Duarte CB: Role of the brain-derived neurotrophic factor at glutamatergic synapses. British journal of pharmacology 2008, 153 Suppl 1:S310–324.

18. Afonso P, De Luca P, Carvalho RS, Cortes L, Pinheiro P, Oliveiros B, Almeida RD, Mele M, Duarte CB: BDNF increases synaptic NMDA receptor abundance by enhancing the local translation of Pyk2 in cultured hippocampal neurons. Sci Signal 2019, 12:eaav3577.

19. Crozier RA, Black IB, Plummer MR: Blockade of NR2B-containing NMDA receptors prevents BDNF enhancement of glutamatergic transmission in hippocampal neurons. Learning & memory (Cold Spring Harbor, NY) 1999, 6:257–266.

20. Hofer M, Pagliusi SR, Hohn A, Leibrock J, Barde YA: Regional distribution of brain-derived neurotrophic factor mRNA in the adult mouse brain. The EMBO journal 1990, 9:2459–2464.

21. Altar CA, Cai N, Bliven T, Juhasz M, Conner JM, Acheson AL, Lindsay RM, Wiegand SJ: Anterograde transport of brain-derived neurotrophic factor and its role in the brain. Nature 1997, 389:856–860.

22. Baquet ZC, Gorski JA, Jones KR: Early striatal dendrite deficits followed by neuron loss with advanced age in the absence of anterograde cortical brain-derived neurotrophic factor. The Journal of neuroscience : the official journal of the Society for Neuroscience 2004, 24:4250–4258.

23. Canals JM, Pineda JR, Torres-Peraza JF, Bosch M, Martín-Ibañez R, Muñoz MT, Mengod G, Ernfors P, Alberch J: Brain-derived neurotrophic factor regulates the onset and severity of motor dysfunction associated with enkephalinergic neuronal degeneration in Huntington’s disease. The Journal of neuroscience : the official journal of the Society for Neuroscience 2004, 24:7727–7739.

24. Zuccato C, Marullo M, Conforti P, MacDonald ME, Tartari M, Cattaneo E: Systematic assessment of BDNF and its receptor levels in human cortices affected by Huntington’s disease. Brain pathology (Zurich, Switzerland) 2008, 18:225–238.

25. Brito V, Puigdellívol M, Giralt A, del Toro D, Alberch J, Ginés S: Imbalance of p75(NTR)/TrkB protein expression in Huntington’s disease: implication for neuroprotective therapies. Cell death & disease 2013, 4:e595.

26. Gharami K, Xie Y, An JJ, Tonegawa S, Xu B: Brain-derived neurotrophic factor over-expression in the forebrain ameliorates Huntington’s disease phenotypes in mice. J Neurochem 2008, 105:369–379.

27. Xie Y, Hayden MR, Xu B: BDNF overexpression in the forebrain rescues Huntington’s disease phenotypes in YAC128 mice. The Journal of neuroscience : the official journal of the Society for Neuroscience 2010, 30:14708–14718.

28. Nguyen KQ, Rymar VV, Sadikot AF: Impaired TrkB Signaling Underlies Reduced BDNF-Mediated Trophic Support of Striatal Neurons in the R6/2 Mouse Model of Huntington’s Disease. Frontiers in cellular neuroscience 2016, 10:37.

29. Li JL, Hayden MR, Almqvist EW, Brinkman RR, Durr A, Dodé C, Morrison PJ, Suchowersky O, Ross CA, Margolis RL, et al: A genome scan for modifiers of age at onset in Huntington disease: The HD MAPS study. American journal of human genetics 2003, 73:682–687.

30. Wexler NS, Lorimer J, Porter J, Gomez F, Moskowitz C, Shackell E, Marder K, Penchaszadeh G, Roberts SA, Gayán J, et al: Venezuelan kindreds reveal that genetic and environmental factors modulate Huntington’s disease age of onset. Proceedings of the National Academy of Sciences of the United States of America 2004, 101:3498–3503.

31. Identification of Genetic Factors that Modify Clinical Onset of Huntington’s Disease. Cell 2015, 162:516–526.

32. Chaves G, Stanley J, Pourmand N: Mutant Huntingtin Affects Diabetes and Alzheimer’s Markers in Human and Cell Models of Huntington’s Disease. Cells 2019, 8.

33. Glerup S, Bolcho U, Molgaard S, Boggild S, Vaegter CB, Smith AH, Nieto-Gonzalez JL, Ovesen PL, Pedersen LF, Fjorback AN, et al: SorCS2 is required for BDNF-dependent plasticity in the hippocampus. Mol Psychiatry 2016, 21:1740–1751.

34. Glerup S, Olsen D, Vaegter CB, Gustafsen C, Sjoegaard SS, Hermey G, Kjolby M, Molgaard S, Ulrichsen M, Boggild S, et al: SorCS2 Regulates Dopaminergic Wiring and Is Processed into an Apoptotic Two-Chain Receptor in Peripheral Glia. Neuron 2014, 82:1074–1087.

35. Olsen D, Wellner N, Kaas M, de Jong IEM, Sotty F, Didriksen M, Glerup S, Nykjaer A: Altered dopaminergic firing pattern and novelty response underlie ADHD-like behavior of SorCS2-deficient mice. Transl Psychiatry 2021, 11:74.

36. Malik AR, Willnow TE: VPS10P Domain Receptors: Sorting Out Brain Health and Disease. Trends Neurosci 2020, 43:870–885.

37. Glerup S, Olsen D, Vaegter CB, Gustafsen C, Sjoegaard SS, Hermey G, Kjolby M, Molgaard S, Ulrichsen M, Boggild S, et al: SorCS2 regulates dopaminergic wiring and is processed into an apoptotic two-chain receptor in peripheral glia. Neuron 2014, 82:1074–1087.

38. Mangiarini L, Sathasivam K, Seller M, Cozens B, Harper A, Hetherington C, Lawton M, Trottier Y, Lehrach H, Davies SW, Bates GP: Exon 1 of the HD gene with an expanded CAG repeat is sufficient to cause a progressive neurological phenotype in transgenic mice. Cell 1996, 87:493–506.

39. Tomé S, Manley K, Simard JP, Clark GW, Slean MM, Swami M, Shelbourne PF, Tillier ER, Monckton DG, Messer A, Pearson CE: MSH3 polymorphisms and protein levels affect CAG repeat instability in Huntington’s disease mice. PLoS genetics 2013, 9:e1003280.

40. Hawes SL, Gillani F, Evans RC, Benkert EA, Blackwell KT: Sensitivity to theta-burst timing permits LTP in dorsal striatal adult brain slice. Journal of neurophysiology 2013, 110:2027–2036.

41. López-Benito S, Sánchez-Sánchez J, Brito V, Calvo L, Lisa S, Torres-Valle M, Palko ME, Vicente-García C, Fernández-Fernández S, Bolaños JP, et al: Regulation of BDNF Release by ARMS/Kidins220 through Modulation of Synaptotagmin-IV Levels. The Journal of neuroscience : the official journal of the Society for Neuroscience 2018, 38:5415–5428.

42. Sahu MP, Nikkilä O, Lågas S, Kolehmainen S, Castrén E: Culturing primary neurons from rat hippocampus and cortex. Neuronal Signal 2019, 3:Ns20180207.

43. Fred SM, Laukkanen L, Brunello CA, Vesa L, Göös H, Cardon I, Moliner R, Maritzen T, Varjosalo M, Casarotto PC, Castrén E: Pharmacologically diverse antidepressants facilitate TRKB receptor activation by disrupting its interaction with the endocytic adaptor complex AP-2. The Journal of biological chemistry 2019, 294:18150–18161.

44. Rezgaoui M, Hermey G, Riedel IB, Hampe W, Schaller HC, Hermans-Borgmeyer I: Identification of SorCS2, a novel member of the VPS10 domain containing receptor family, prominently expressed in the developing mouse brain. Mech Dev 2001, 100:335–338.

45. Lovinger DM: Neurotransmitter roles in synaptic modulation, plasticity and learning in the dorsal striatum. Neuropharmacology 2010, 58:951–961.

46. Gerfen CR, Surmeier DJ: Modulation of striatal projection systems by dopamine. Annu Rev Neurosci 2011, 34:441–466.

47. Ehrlich ME: Huntington’s disease and the striatal medium spiny neuron: cell-autonomous and non-cell-autonomous mechanisms of disease. Neurotherapeutics 2012, 9:270–284.

48. Ginés S, Bosch M, Marco S, Gavaldà N, Díaz-Hernández M, Lucas JJ, Canals JM, Alberch J: Reduced expression of the TrkB receptor in Huntington’s disease mouse models and in human brain. The European journal of neuroscience 2006, 23:649–658.

49. Knüsel B, Rabin SJ, Hefti F, Kaplan DR: Regulated neurotrophin receptor responsiveness during neuronal migrationand early differentiation. The Journal of neuroscience : the official journal of the Society for Neuroscience 1994, 14:1542–1554.

50. Di Lieto A, Rantamäki T, Vesa L, Yanpallewar S, Antila H, Lindholm J, Rios M, Tessarollo L, Castrén E: The responsiveness of TrkB to BDNF and antidepressant drugs is differentially regulated during mouse development. PloS one 2012, 7:e32869.

51. Guo W, Nagappan G, Lu B: Differential effects of transient and sustained activation of BDNF-TrkB signaling. Developmental neurobiology 2018, 78:647–659.

52. Volianskis A, France G, Jensen MS, Bortolotto ZA, Jane DE, Collingridge GL: Long-term potentiation and the role of N-methyl-D-aspartate receptors. Brain research 2015, 1621:5–16.

53. Lin SY, Wu K, Levine ES, Mount HT, Suen PC, Black IB: BDNF acutely increases tyrosine phosphorylation of the NMDA receptor subunit 2B in cortical and hippocampal postsynaptic densities. Brain research Molecular brain research 1998, 55:20–27.

54. Xu F, Plummer MR, Len GW, Nakazawa T, Yamamoto T, Black IB, Wu K: Brain-derived neurotrophic factor rapidly increases NMDA receptor channel activity through Fyn-mediated phosphorylation. Brain research 2006, 1121:22–34.

55. Otis JM, Fitzgerald MK, Mueller D: Infralimbic BDNF/TrkB enhancement of GluN2B currents facilitates extinction of a cocaine-conditioned place preference. The Journal of neuroscience : the official journal of the Society for Neuroscience 2014, 34:6057–6064.

56. Smith-Dijak AI, Sepers MD, Raymond LA: Alterations in synaptic function and plasticity in Huntington disease. J Neurochem 2019, 150:346–365.

57. Mitre M, Mariga A, Chao MV: Neurotrophin signalling: novel insights into mechanisms and pathophysiology. Clinical science (London, England : 1979) 2017, 131:13–23.

58. Milnerwood AJ, Raymond LA: Early synaptic pathophysiology in neurodegeneration: insights from Huntington’s disease. Trends Neurosci 2010, 33:513–523.

59. Puigdellívol M, Cherubini M, Brito V, Giralt A, Suelves N, Ballesteros J, Zamora-Moratalla A, Martín ED, Eipper BA, Alberch J, Ginés S: A role for Kalirin-7 in corticostriatal synaptic dysfunction in Huntington’s disease. Hum Mol Genet 2015, 24:7265–7285.

60. Jia Y, Gall CM, Lynch G: Presynaptic BDNF promotes postsynaptic long-term potentiation in the dorsal striatum. The Journal of neuroscience : the official journal of the Society for Neuroscience 2010, 30:14440–14445.

61. Li W, Pozzo-Miller L: Differences in GluN2B-Containing NMDA Receptors Result in Distinct Long-Term Plasticity at Ipsilateral versus Contralateral Cortico-Striatal Synapses. eNeuro 2019, 6.

62. Suen PC, Wu K, Levine ES, Mount HT, Xu JL, Lin SY, Black IB: Brain-derived neurotrophic factor rapidly enhances phosphorylation of the postsynaptic N-methyl-D-aspartate receptor subunit 1. Proceedings of the National Academy of Sciences of the United States of America 1997, 94:8191–8195.

63. Mizuno M, Yamada K, He J, Nakajima A, Nabeshima T: Involvement of BDNF receptor TrkB in spatial memory formation. *Learning & memory (Cold Spring Harbor*, NY) 2003, 10:108–115.

64. Ma Q, Yang J, Milner TA, Vonsattel JG, Palko ME, Tessarollo L, Hempstead BL: SorCS2-mediated NR2A trafficking regulates motor deficits in Huntington’s disease. JCI insight 2017, 2.

65. Ayton S, Lei P, Appukuttan AT, Renoir T, Foliaki S, Chen F, Adlard PA, Hannan AJ, Bush AI: Brain Zinc Deficiency Exacerbates Cognitive Decline in the R6/1 Model of Huntington’s Disease. Neurotherapeutics 2020, 17:243–251.

66. Yang J, Ma Q, Dincheva I, Giza J, Jing D, Marinic T, Milner TA, Rajadhyaksha A, Lee FS, Hempstead BL: SorCS2 is required for social memory and trafficking of the NMDA receptor. Mol Psychiatry 2021, 26:927–940.

67. Morella I, Hallum H, Brambilla R: Dopamine D1 and Glutamate Receptors Co-operate With Brain-Derived Neurotrophic Factor (BDNF) and TrkB to Modulate ERK Signaling in Adult Striatal Slices. Frontiers in cellular neuroscience 2020, 14:564106.

68. Yang K, Trepanier C, Sidhu B, Xie YF, Li H, Lei G, Salter MW, Orser BA, Nakazawa T, Yamamoto T, et al: Metaplasticity gated through differential regulation of GluN2A versus GluN2B receptors by Src family kinases. The EMBO journal 2012, 31:805–816.

69. Koch ET, Raymond LA: Dysfunctional striatal dopamine signaling in Huntington’s disease. Journal of neuroscience research 2019, 97:1636–1654.

70. Visanji NP, Kamali Sarvestani I, Creed MC, Shams Shoaei Z, Nobrega JN, Hamani C, Hazrati LN: Deep brain stimulation of the subthalamic nucleus preferentially alters the translational profile of striatopallidal neurons in an animal model of Parkinson’s disease. Frontiers in cellular neuroscience 2015, 9:221.

71. Andrew SE, Goldberg YP, Kremer B, Telenius H, Theilmann J, Adam S, Starr E, Squitieri F, Lin B, Kalchman MA, et al.: The relationship between trinucleotide (CAG) repeat length and clinical features of Huntington’s disease. Nature genetics 1993, 4:398–403.

72. Djoussé L, Knowlton B, Hayden M, Almqvist EW, Brinkman R, Ross C, Margolis R, Rosenblatt A, Durr A, Dode C, et al: Interaction of normal and expanded CAG repeat sizes influences age at onset of Huntington disease. American journal of medical genetics Part A 2003, 119a:279–282.

73. Labadorf A, Hoss AG, Lagomarsino V, Latourelle JC, Hadzi TC, Bregu J, MacDonald ME, Gusella JF, Chen JF, Akbarian S, et al: RNA Sequence Analysis of Human Huntington Disease Brain Reveals an Extensive Increase in Inflammatory and Developmental Gene Expression. PloS one 2015, 10:e0143563.

